# Immune interaction signatures in adipose tissue fibroblasts in obesity-associated atherosclerosis

**DOI:** 10.1101/2025.09.02.673686

**Authors:** Mikkola Lea, Bölük Aydin, Piipponen Minna, Mikocziova Ivana, Valkonen Mira, Fagersund Jimmy, Hakovirta Harri, Saraste Antti, Hernández de Sande Ana, Palani Senthil, Örd Tiit, Roivainen Anne, Ruusuvuori Pekka, Heinäniemi Merja, Kaikkonen U. Minna, Lönnberg Tapio

## Abstract

Atherosclerosis involves changes in the vascular wall and surrounding perivascular adipose tissue, yet the cellular contributors to disease progression remain incompletely understood. Obesity exacerbates atherogenesis, but the cell types driving this aggravation are unclear. We aimed to define the key cell populations across tissues in a highly atherogenic mouse model under obese and normal-weight conditions and to identify obesity-associated cellular changes.

We employed 5’ single-cell RNA sequencing combined with antibody staining in Ldlr^-/-^Apob^100/100^ male mice fed either a high-fat or control diet. Aorta, perivascular and epididymal adipose tissues, and spleen were analyzed, with CD45⁺ enrichment of aortic samples and CITE-seq using a 138-antibody panel. Key findings were validated in mice by immunohistochemistry and multiplexed immunofluorescence and explored in human aorta and carotid arteries using spatial transcriptomics.

Analysis of ∼46,000 cells enabled characterization of cell states, gene enrichment, regulon activity, and inferred interactions. Adipose-derived fibroblast subsets displayed immune-associated transcriptional programs in obesity. *Pi16⁺* progenitor fibroblasts were reduced alongside marked PVAT remodeling, and the top mouse differentially expressed genes exhibited clear spatial patterning in human arteries.

## Introduction

Atherosclerosis is a pervasive common complex disorder that is greatly impacted by genetic components. Large genome-wide association studies have identified hundreds of risk loci but their predictive power and especially understanding of the disease mechanisms is still limited^1^. Single-cell and spatial omics (e.g. transcriptomics, proteomics and epigenetics) studies have expanded in recent years to enable high-resolution profiling of cellular heterogeneity, in-depth investigation of transcriptional cell regulation, and cell-cell and cell-tissue microenvironment (TME) crosstalk in health and disease, including atherosclerosis^2^. Immune-structural cell interactions have been recognized to be pivotal for the maintenance of tissue homeostasis in healthy tissues and during disease states^3,4^. Especially fibroblasts have been under growing interest in atherosclerosis studies in the recent years ^5^. Activated adventitial fibroblasts have been indicated as having multifaceted functions during atherosclerosis progression, for instance in proinflammatory signalling, extracellular matrix modulation, and immune cell recruitment via chemotaxis^5^. Switching off inflammatory programs is important after resolving an acute inflammation, and if this fails, chronic inflammation may occur. Fibroblasts that have lost their ability to switch away from the inflammatory program could be at the focal point of persistent inflammation^6^.

Instead of concentrating only on immune cells and/or only on arterial tissues, our objectives were to map the immune-stromal landscape of multiple tissues and investigate the intricate cell interactions in an atherosclerosis mouse model to better understand the drivers of the disease during obesity. To this end, we generated data that included a wide variety of immune and non-immune cell types and multiple tissue types that might harbor cells crucial to the development of atherosclerosis. In addition to the whole aorta, we collected perivascular adipose tissue (PVAT), epididymal white adipose tissue (eWAT), and the whole spleen. Spleen was included due to its important role in lipid metabolism, homing and transforming of lymphocytes, macrophage activation, and potentially substantial participation in the atherosclerosis-associated immune responses ^7,8^. Meanwhile, PVAT and eWAT were important target tissues because obesity has a major impact on the development of atherosclerosis, and it has been shown that both the white adipose tissue (WAT), especially in the visceral depots, and PVAT may convert to a proinflammatory phenotype during obesity induced by caloric intake excess^9,10^. Our data included samples from both obese and non-obese mice (Fig 1A) that had been fed either a high-calorie or normal chow diet for three months, thus modelling the effect of caloric intake excess on the development of atherosclerosis.

**Figure 1.**
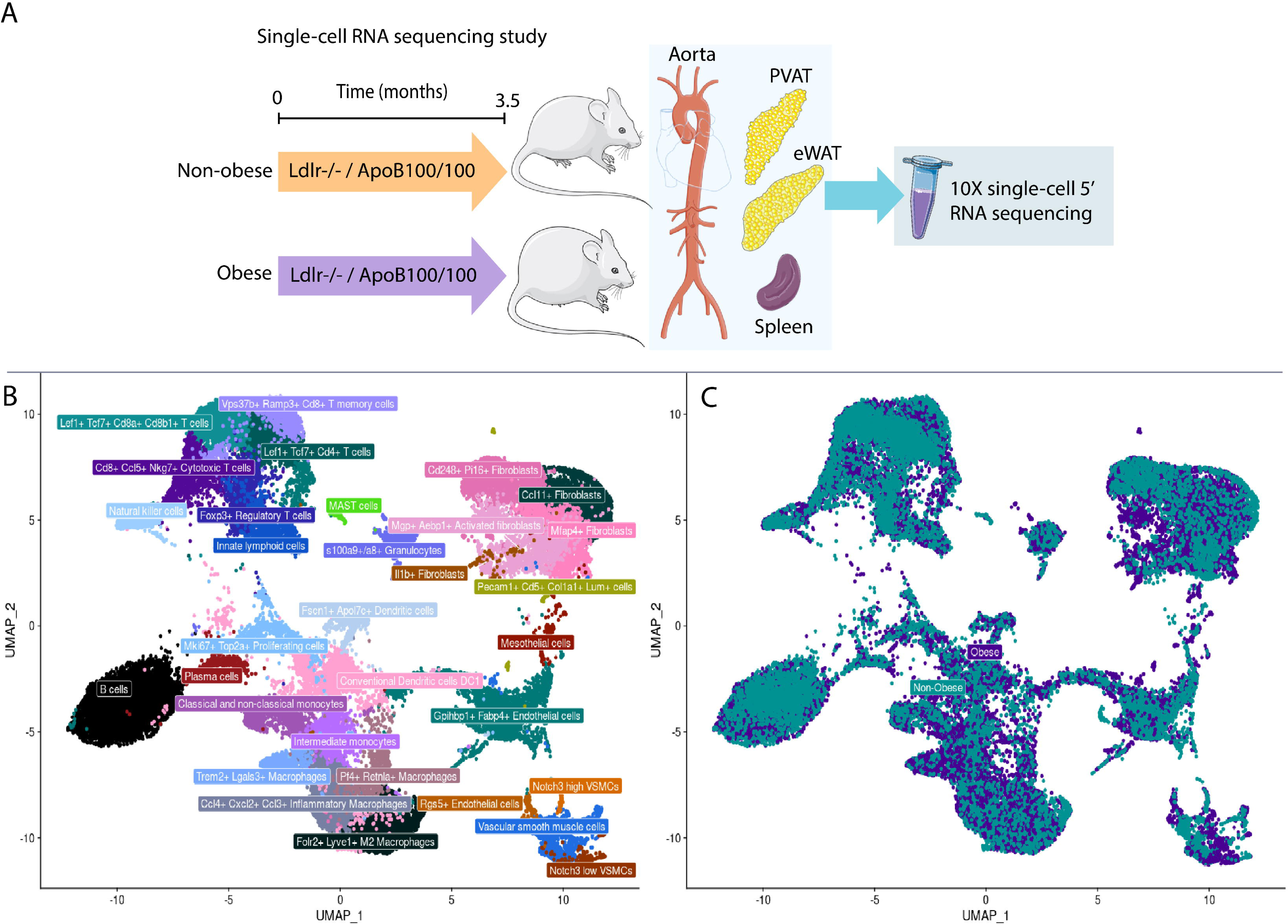
Study design, cell type annotations, and obese/non-obese status of cells. A) Study design of the single-cell RNA sequencing study (Image created using Servier Medical Art, licensed under CC BY 4.0), B) UMAP projection of the integrated data with cell type annotations, and C) UMAP projection of the integrated data showing the obesity status of each cell. Number of cells is 45940.

We have constructed a uniquely diverse multi-tissue single-cell data of atherosclerosis murine models representing the disease states during obesity and normal body weight, which enabled the mapping of the most pivotal cell types and changes in their gene expression profiles, as well as study of their regulons and cell interactions. We found that PVAT- and eWAT-derived fibroblasts are drastically impacted by obesity in this atherosclerosis model. Moreover, we showed that in addition to strong potential in interactions with immune cells, some of these fibroblasts demonstrated marked immune-regulation potential that has not been described previously in the context of atherosclerosis and obesity.

## Methods

The detailed methods, including information about the reagents, antibodies, analysis parameters, data processing steps, information about the programs/R-packages, as well as links to scripts, protocols, and used materials to all the workflows are provided in Supplementary Methods.

### Data availability

The single-cell RNA sequencing data is available in the Gene Expression Omnibus database (GSE235895). Due to data sensitivity, the spatial transcriptomics data of human samples is not available via open repositories, but the non-sensitive count matrices can be shared to trusted parties on request from VarHa (wellbeing services county of Southwest Finland) research centre.

IHC-based cell counting data of PI16+ fibroblasts is available at <u>10.6084/m9.figshare.28956140</u>, and the data for the morphometric analysis at <u>10.6084/m9.figshare.28956104</u>. Original images of CD19 and CD74 antibody staining on mouse PVAT-aorta are available at <u>10.6084/m9.figshare.29279045</u> (CD19) and <u>10.6084/m9.figshare.30030004</u> (CD74). Human aorta and carotid artery H&E, CD68 and CD206 staining images are available at <u>10.6084/m9.figshare.30016450</u>.

### Animals

Animal experiments were approved by the National Experiment Animal Board of Finland (permission numbers ESAVI/4567/2018, ESAVI/6772/2018, and ESAVI/17197/2021) and were carried out in accordance with the Finnish Act on Animal Experimentation and EU Directive 2010/EU/63 on the protection of animals used for scientific purposes.

### Sample collection and processing for single-cell transcriptomics

Ten male Ldlr⁻/⁻ ApoB100/100 mice (C57BL/6J) were fed either a high-fat diet (n=5) or regular chow (n=5) for 3.5 months. Mice were age-matched and sacrificed under isoflurane anesthesia (induction 4-5 %, maintenance 2-2.5 %), followed by cardiac puncture and cervical dislocation. Perfusion was performed with 10 ml ice-cold DPBS containing 10 U/ml heparin.

Aorta (from ascending to renal arteries), PVAT, eWAT, and spleen were collected post-perfusion. PVAT was separated from aortas on a cold block and tissues were processed into single-cell suspensions using a published protocol (link in Supplementary Methods). RBC lysis was performed, and cell viability assessed using acridine orange/propidium iodide staining. CD45⁺ enrichment was performed on pooled aorta samples using Miltenyi MicroBeads. To retain CD45⁻ cells, 30 µl of the original sample was mixed back in (20–25% original: 75–80% enriched). Cells were stained with a 138-antibody TotalSeq-C panel by BioLegend, after which we continued with the single-cell immune profiling protocol by 10X Genomics.

For validation studies, ten additional mice were obtained from the University of Eastern Finland. Five were fed high-fat diet and five chow similarly as before. Aorta-PVAT samples were fixed with 4% paraformaldehyde and embedded for immunohistochemistry (IHC) and multiplexed immunofluorescence (mIF) experiments.

### TotalSeq antibody staining of cell surface proteins

Approximately 2.5×10⁵ cells per tissue were stained. Fc receptors were blocked with anti-CD16/CD32 (PN-BE0307, InVivoMAb) in 1% BSA-PBS. The TotalSeq-C cocktail (PN-900002539, BioLegend) was reconstituted per manufacturer’s instructions. Cells were incubated with 12.5 µl antibody panel and 75 µl 1% BSA-PBS on ice for 30 minutes. After two washes (0.04% and 1% BSA-PBS), cells were resuspended at 300–1000 cells/µl in 1% BSA-PBS.

### Single-cell sequencing and preprocessing of the sequencing data

Eight pooled samples (obese and non-obese states) were processed using 10X Genomics Chromium Next GEM Single Cell 5’ v2 kits (details in the Supplementary Methods) following manufacturer’s instructions. Libraries for gene expression and surface proteins were prepared and sequenced at Novogene on NovaSeq6000 (S4 flow cell, 150 bp paired-end). Quality controls (QC) throughout the workflow were performed using Bioanalyzer 2100 (Agilent Technologies, US). Data were pre-processed using Cell Ranger v6 (multi pipeline).

### Quality control and sample integration

Initial sequencing data QC followed by integration was performed in R on a high-performance computing environment of CSC - IT Center for Science (Finland). Ambient RNA was removed using SoupX. Cells with <200 or>5000 features or high mitochondrial content (>5–10%) were excluded. Doublets were removed using DoubletFinder.

The cleaned data was merged, after which filtering was done to remove some technical noise genes (see Supplementary Methods). The data was then renormalized and scaled. One antibody was removed from the CITE-seq data due to a quality issue reported by the manufacturer (BioLegend). CITE-seq data was then normalized using centered log-ratio and denoised using the DSB package with isotype controls. Each sample was processed and integrated separately. Principal component analysis and subsequent UMAP-dimensional reduction were performed, and Harmony was used for batch correction. The appropriate clustering resolution was determined using Clustree and visual inspection. A multipositive cluster expressing both B and T cell markers was excluded, reducing the final cell count from 47,495 to 45,940.

### Cell annotation and differential gene expression analysis

Cell types were annotated using SingleR and celldex (MouseRNAseqData reference) and manual inspection of CITE-seq and gene expression markers. Differential expression was assessed using Wilcoxon rank sum test with adjusted p < 0.05.

### Gene set enrichment analysis

GSEA was performed using the web-based g:Profiler (GO, KEGG, Reactome databases). Immune-related terms (listed in Supplementary Methods) were extracted using a custom R script (available) and visualized with enrichplot and multienrichjam.

### Trajectory analysis

Trajectory inference was performed using R-package Totem on SingleCellExperiment objects. Seurat objects were converted using SingleCellExperiment, and dimensionality reduction was performed using dyndimred.

### Single-cell regulatory network inference and clustering (SCENIC) analysis

SCENIC analysis (pySCENIC Python-package) was used to infer transcription factor activity via GRNBoost2, cisTarget, and AUCell. Regulon specificity scores were calculated using Jensen–Shannon divergence.

### Cell interaction analysis

NicheNet was used to infer ligand–target interactions between defined sender cell types (e.g., fibroblast subsets, macrophages, T cells, endothelial cells) in PVAT. The Mki67 and Top2a positive cell cluster was excluded from this analysis, because it can represent multiple cell types. Input matrices were obtained from Zenodo (see Supplementary Methods).

### Sample collection of human carotid arteries and aorta for spatial transcriptomics

Human aorta and carotid artery samples were collected from three patients (details in Supplementary Methods) during aorto-femoral bypass surgery due to chronic, symptomatic peripheral artery disease. Patients gave written informed consent, and the study conforms to the Declaration of Helsinki. The institutional review boards of the Hospital District of Southwest Finland and Turku University Hospital approved the study (permit number T03/024/19, ethical statement number T75_2011). Sections were stained with H&E and IF to guide region selection.

### Histological and immunofluorescence staining of human aorta and carotid artery sections

Tissues were embedded in optimal cutting temperature compound, frozen at −70°C, and sectioned at 6 µm. H&E staining was performed and slides scanned. Adjacent sections were formalin-fixed, treated with epitope retrieval, and blocked. Double staining was performed using antibodies against CD68, Mannose receptor and Folate receptor-beta (CD206), followed by Alexa Fluor-conjugated secondary antibodies. Slides were mounted and scanned using a fluorescence scanner. Regions of interest were selected for spatial transcriptomics experiment (Supplementary Fig 1).

### Spatial transcriptomics sample processing and sequencing

Spatial transcriptomics was performed using 10X Genomics Visium (V1). Two technical replicates per sample were generated (total eight sections). Imaging was done using Nikon Eclipse Ti2-E. RNA integrity and permeabilization optimization were assessed. Libraries were sequenced on NovaSeq SP flow cell and processed with Space Ranger v1 by the Finnish Functional Genomics Centre. Data QC was performed in Seurat. Low-quality spots were filtered based on gene count and mitochondrial/ribosomal content thresholds. Data normalization, scaling, and finding of variable features before DE analysis were done similar to the single-cell workflow.

### Immunohistochemistry and multiplexed immunofluorescence staining of mouse tissues

Aorta-PVAT samples were paraffin-embedded and stained at the Histology core facility, University of Turku. Routine IHC included antigen retrieval, semiautomated staining, blocking, primary/secondary antibody incubation, DAB substrate incubation, and hematoxylin counterstaining. Multiplex-IF of PI16, F4/80, CD4, CD3 was performed using automated Leica BOND RX with four staining rounds including: protein blocking, primary/secondary antibody incubations, tyramide signal amplification, and antibody retrieval between rounds. After mounting imaging was done with Leica Thunder microscope at the Cell Imaging and Cytometry core facility, at Turku Bioscience Centre.

### Tissue imaging and image analyses (PI16⁺ cell counts and morphometric analysis)

For the PI16+ cell counting and morphometric analysis imaging was performed using Nikon Eclipse Ti2-E and Pannoramic P1000. Image annotation for the cell counting was done with QuPath. Images and their metadata, which were used in the analyses is available (see Data availability). Morphometric analysis was conducted using Visiopharm software with deep learning classifiers and Visiopharm’s adapted adipose tissue workflows at the Medisiina Imaging Centre, University of Turku. The generated data was analyzed in R using stats and dplyr, and visualized with ggplot2. Statistical tests included Shapiro-Wilk for normality and one-way Welch’s t-test.

PI16⁺ fibroblast counts were estimated using a domain-adapted nucleus detection method. Two-step domain adaptation was applied using pseudo labels and retraining using randomly chosen detections. Final detection used thresholding of the brown channel. Subsequent statistical analysis workflow first examined group differences by fitting a generalized linear regression model using beta distribution and then evaluated the model fit. Next, the estimated mean cell counts were compared between groups with Type III ANOVA. Finally, estimated marginal means were generated to inspect how the estimated group means diverged.

## Results

### Tissue-specific characterization of the cellular landscape in murine model of atherosclerosis

Using 5’ single-cell RNA sequencing combined with CITE-seq (Fig 1A), we generated a data of ∼46000 cells. The unsupervised clustering of the integrated data which included all the tissues and both obesity states identified 40 clusters. We defined 13 major cell types that corresponded to a total of 32 different cell states and were clearly present in both model states (Fig 1B and C). These included five different T cell populations (T memory cells, *Cd8*+ and *Cd4*+ T cells positive for *Lef1* and *Tcf7*, *Cd8*+ cytotoxic T cells, and regulatory T cells), five fibroblast (*Pi16*+, *Ccl11*+, *Mfap*+, *Mgp*+, and *Il1b*+) subpopulations, four macrophage populations (M2, M1, *Trem2*+ foamy, and *Pf4*+ *Retnla*+ macrophages), three vascular smooth muscle cell (VSMC) populations, two B cell (B cells and plasma cells), two dendritic cell (conventional and *Fscn1*+ *Apol7c*+ DC), two endothelial cell, two monocyte (intermediate and classical and non-classical as one), one granulocyte, one innate lymphoid cell (ILC), one natural killer (NK), one mast cell, and one mesothelial cell population. Moreover, one mixed *Mki67*+ *Top2*a+ proliferating population, and a separate small cluster demonstrating high expression of both endothelial and fibroblast markers (*Pecam1*, *Cdh5*, *Col1a1*, and *Lum*) were identified. The top five most abundant cell types per tissue and obesity state are shown in Table 1.

**Table 1.** Top five most abundant cell types in obese and non-obese samples per tissue.

| Tissue | Cell type in obese mice | Proportion of the total cell count in the sample | Cell type in non-obese mice | Proportion of the total cell count in the sample |
| --- | --- | --- | --- | --- |
| Aorta | B cells | 23.2 (614/2651) | B cells | 24.7 (1652/6684) |
|  | VSMC | 8.4 (223/2651) | Conventional Dendritic cells DC1 | 8.9 (594/6684) |
|  | *Vps37b+ Ramp3+ Cd8+* T memory cells | 7.7 (203/2651) | *Ccl4+ Cxcl2+ Ccl3+* Inflammatory Macrophages | 8.7 (584/6684) |
|  | *Ccl4+ Cxcl2+ Ccl3+* Inflammatory Macrophages | 7.6 (201/2651) | *Vps37b+ Ramp3+ Cd8+* T memory cells | 8.6 (575/6684) |
|  | *Trem2+ Lgals3+* Macrophages | 7.1 (187/2651) | *Folr2+ Lyve1+* M2 Macrophages | 8.1 (542/6684) |
| eWAT | *Gpihbp1+ Fabp4+* Endothelial cells | 16.5 (1314/7986) | *Gpihbp1+ Fabp4+* Endothelial cells | 19.3 (1238/6411) |
|  | *Folr2+ Lyve1+* M2 Macrophages | 14.4 (1152/7986) | *Folr2+ Lyve1+* M2 Macrophages | 15.1 (970/6411) |
|  | *Ccl11+* Fibroblasts | 13.1 (1043/7986) | *Ccl11+* Fibroblasts | 10.4 (668/6411) |
|  | *Ccl4+ Cxcl2+ Ccl3+* Inflammatory Macrophages | 6.6 (530/7986) | *Cd248+ Pi16+* Fibroblasts | 7.5 (482/6411) |
|  | *Cd248+ Pi16+* Fibroblasts | 6.0 (478/7986) | *Ccl4+ Cxcl2+ Ccl3+* Inflammatory Macrophages | 7.4 (477/6411) |
| PVAT | *Mgp+ Aebp1+* Activated fibroblasts | 11.5 (843/7343) | B cells | 19.5 (1050/5377) |
|  | *Mfap4+* Fibroblasts | 11.4 (835/7343) | *Cd248+ Pi16+* Fibroblasts | 11.6 (623/5377) |
|  | *Cd248+ Pi16+* Fibroblasts | 8.6 (632/7343) | *Ccl11+* Fibroblasts | 7.7 (412/5377) |
|  | *Ccl11+* Fibroblasts | 8.4 (618/7343) | *Mfap4+* Fibroblasts | 7.2 (385/5377) |
|  | *Gpihbp1+ Fabp4+* Endothelial cells | 6.7 (492/7343) | Classical and non-classical monocytes | 6.1 (328/5377) |
| Spleen | B cells | 20.6 (869/4228) | B cells | 26.4 (1387/5260) |
|  | *Lef1+ Tcf7+ Cd4+* T cells | 20.5 (868/4228) | *Lef1+ Tcf7+ Cd4+* T cells | 21.1 (1108/5260) |
|  | *Lef1+ Tcf7+ Cd8a+ Cd8b1+* T cells | 18.2 (769/4228) | *Lef1+ Tcf7+ Cd8a+ Cd8b1+* T cells | 19.8 (1040/5260) |
|  | *Foxp3+* Regulatory T cells | 9.7 (411/4228) | *Foxp3+* Regulatory T cells | 9.2 (485/5260) |
|  | *Cd8+ Ccl5+ Nkg7+* Cytotoxic T cells | 8.7 (366/4228) | *Cd8+ Ccl5+ Nkg7+* Cytotoxic T cells | 7.8 (410/5260) |

The cells from the aortas showed expected enrichment of different immune cell populations (Table 1). However, surprisingly, the most abundant cell type in aorta (as well as in PVAT) were B cells, which was not expected based on literature, and we hypothesized that lymph node-derived B cell contamination was likely impacting the cell counts in these tissues. To confirm this, we carried out a CD19 IHC staining in the validation set of aorta-PVAT sections. The marker was detected in the lymph nodes and lymph vessels but not elsewhere in the PVAT or aorta (Supplementary Fig 2), confirming their likely origin also in the single-cell data.

### Differential gene expression analysis highlights Cd74 and Apoe as key genes between the obesity states in multiple cell types across tissues

We performed differential gene expression analyses between the obese and non-obese atherosclerosis states in key cell types. We report changes in cell populations with at least one gene showing Log2 Fold change (Log2FC) greater than 1.5 (full lists of DE genes are provided in Supplementary Tables 2-4).

The most prominent gene expression changes were observed in the two adipose tissues, PVAT and eWAT (Fig 2A and B). In spleen none of the observed gene expression changes reached the set Log2FC threshold, while in aorta, the changes were generally of smaller effect and were restricted to *Cd8+* Tmem cells and *Fscn1+ Apol7c+* DC populations (Supplementary Fig 3A and B). Moreover, we unexpectedly observed many immunoglobulin variable genes and Joining Chain Of Multimeric IgA And IgM (*Jchain*) among the most upregulated genes in these aorta-derived cells, possibly representing a technical artefact. Unlike the other tissue types, the aortas were enriched for CD45+ cells before pooling, which could have introduced more debris (see Methods). The removal of ambient RNA, even when focusing on these immunoglobulin transcripts specifically, as recommended in^11^ where a similar phenomenon was observed, did not resolve the issue. To avoid skewing the results by removing these transcripts, we concentrated on the other upregulated genes instead. The most highly upregulated gene in the aorta-derived T memory cells was *Cd74* (CD74 Molecule, also known as HLA-DR antigens-associated invariant chain; Log2FC ∼1.8, Supplementary Table 2), which was also observed upregulated in other cell types. Some histocompatibility 2 antigens (*H2-Aa* and *H2-Eb1*) were also upregulated, but their Log2FCs were less than 0.9 (Supplementary Table 2). In the aortic DC population, the most upregulated genes were *Cxcl2* (C-X-C Motif Chemokine Ligand 2, Log2FC ∼ 1.5), and *Apoe* (Apolipoprotein E, Log2FC ∼ 1.1) (Supplementary Table 2).

**Figure 2.**
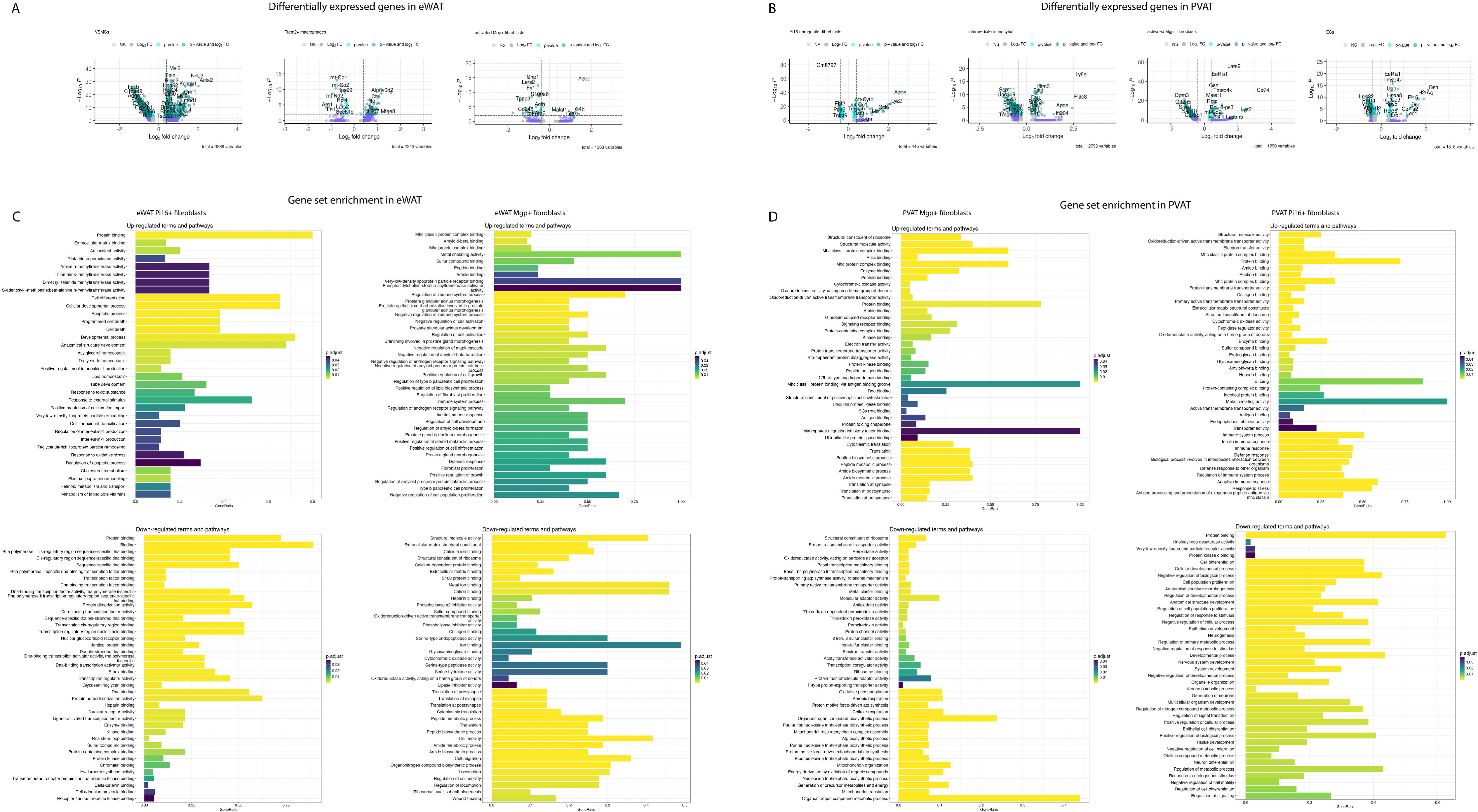
Differentially expressed genes and top hits from the GSEA. Differentially expressed genes between the obese and non-obese states in A) eWAT VSMCs (N=582), *Trem2+* macrophages (N=383), and *Mgp+* fibroblasts (N=479), and B) PVAT *Pi16+* fibroblasts (1255), intermediate monocytes (N=276), *Mgp+* fibroblasts (N=1054), and ECs (N=811). Positive fold changes indicate higher expression in the obese state. Statistical test was Wilcoxon rank sum test. Threshold for log2 fold change was 0.5 and threshold for significance was adjusted P-value < 0.05. Top 40 enriched terms based on the up-and downregulated DE genes in eWAT-(C) and PVAT-derived (D) *Pi16+* and *Mgp+* fibroblasts. On the x-axis the gene ratio indicates the number of overlapping genes divided by the total number of reference genes in the gene ontology category. g:Profiler uses Fisher’s one-tailed test for GSEA, and we used adjusted P-value of 0.05 as the threshold for significance (program default g:SCS algorithm, corresponding to an experiment-wide threshold of a=0.05).

The two adipose tissues showed more marked changes between the two atherosclerosis model states. In PVAT activated (*Mgp+*) fibroblasts and *Pi16+* progenitor fibroblasts, one EC population, and intermediate monocytes (Fig 2B) showed notable changes. The highest Log2FCs in the activated fibroblasts were observed for *Cd74*, *Lyz2* (∼ 2.3-3.5, Supplementary Table 3), and three Histocompatibility 2, class II antigens (*H2-Eb1*, *H2-Ab1*, and *H2-Aa,* Log2FC∼2.3-2.5). *Lyz2* and *Cd74* were also upregulated (Log2FC ∼ 2.3-2.4) in the *Pi16+* progenitor fibroblasts, while the other top genes were *Apoe* (Log2FC ∼ 2.6, and the complement chain encoding genes *C1qb*, and *C1qa* (Log2FC ∼ 2.1, Supplementary Table 3). In the ECs, *Cd74* was observed once again with a Log2FC of ∼ 2.1, while some of the other highly expressed genes during obesity were *Gsn* (Gelsolin, Log2FC ∼ 2.3) and the same histocompatibility antigens as in the activated fibroblasts (Log2FC of ∼1.7 – ∼ 2.0). Finally, in the intermediate monocytes, the most upregulated genes in obesity were *Ly6e* (Lymphocyte Antigen 6 Family Member E, Log2FC ∼ 2.9), *Plac8* (Placenta Associated 8, Log2FC ∼ 2.9), *Ly6c2* (Lymphocyte Antigen 6 Family Member G6C, Log2FC ∼ 2.4), and *Apoe* (Log2FC ∼ 1.9).

In eWAT the most remarkable gene expression changes were observed in activated *Mgp+* fibroblasts, *Trem2+* macrophages, and VSMCs (Fig 2A). The most upregulated genes in the activated *Mgp+* fibroblasts were *Apoe* (Log2FC ∼ 1.7), *C4b* (Complement component 4b; Log2FC ∼ 1.5) (Supplementary Table 4). In the *Trem2+* macrophages the most upregulated genes were *Gpnmb* (Glycoprotein Nmb; Log2FC ∼ 1.6), and Mfge8 (Milk Fat Globule EGF And Factor V/VIII Domain Containing; Log2FC ∼ 1.5) (Supplementary Table 4). Finally, the four of the most upregulated genes in VSMCs were *Acta2* (Actin Alpha 2, Smooth Muscle; Log2FC ∼ 2.4), *Tagln* (Transgelin; Log2FC ∼ 2.0), *Nrip2* (Nuclear Receptor Interacting Protein 2; Log2FC∼2.0), and *Mustn1* (Musculoskeletal, Embryonic Nuclear Protein 1; Log2FC ∼ 1.8) (Supplementary Table 4).

### Gene set enrichment analysis highlights the upregulation of immune-related processes in PVAT-derived fibroblasts during obesity

As the different fibroblast subpopulations were recurring in the most highly affected cell types in the differential gene expression analyses, we focused our downstream analyses on them. Log2FC-ranked gene lists revealed a multitude of affected processes in the gene set enrichment analysis. The analysis was done for both up-and downregulated genes with g:Profiler^12^, concentrating on the GO molecular functions and biological processes, and the KEGG and REACTOME pathways. Immune-related phrases were identified via text mining. The top up-and downregulated terms are in Fig 2C (eWAT) and D (PVAT).

First, we investigated the PVAT-derived *Mgp+* and *Pi16+* fibroblasts, which exhibited strong differential gene expression between the obesity states (Fig 2D). GSEA analysis of the upregulated DE genes in PVAT-derived *Pi16+* fibroblasts resulted in 301 hits (Supplementary Table 5). In total, ∼35 % (105/301) of the phrases were immune-related, indicating strong immune activation and possible antigen presentation-related functions during obesity. Meanwhile, the analysis of the downregulated genes revealed 82 affected GO terms (Supplementary Table 5). The only immune-related term was a GO Biological Process “Negative regulation of macrophage activation”. In contrast, similar analysis of eWAT-derived *Pi16+* fibroblasts identified 34 hits among upregulated DE genes and 596 hits among downregulated genes (Supplementary Table 6). Only three upregulated terms were immune-related, all linking specifically to interleukin-1 production or its regulation, and the rest were linked to the homeostasis of triglycerides and lipid levels, and lipoprotein remodelling. Meanwhile, approximately 7% (40/588) of the downregulated GO terms were immune-related, reflecting decreased processes in eWAT-derived *Pi16+* fibroblasts during obesity, including leukocyte dynamics, T cell activation, and antigen/cytokine signalling. Interestingly, fat cell differentiation related functions were also downregulated (Supplementary Table 6). These results highlight that eWAT and PVAT-derived *Pi16+* fibroblasts have very different functions during obesity.

In the *Mgp+* fibroblasts in PVAT, we observed 335 terms based on the upregulated DE genes (Supplementary Table 5). Again, immune-related terms and pathways were highly emphasized to be upregulated; ∼ 37 % (106/286) GO terms and ∼ 43 % (12/28) of the KEGG pathways were related to immunity. Antigen processing and presentation, especially via the MHC class II pathways, were highly represented in the results, as was for *Pi16+* fibroblasts in the same adipose tissue. The GSE-analysis results based on the downregulated DE genes in *Mgp+* fibroblasts in PVAT did not include anything immune-related but were focused on e.g. thermogenesis, cellular oxidative processes and respiration, and detoxification and responses to oxidative stress (Supplementary Table 5). In eWAT, the *Mgp+* activated fibroblasts also had upregulated immune-related functions during obesity (Fig 2C), while processes such as structural molecule activity, ECM structural constituents and ECM binding as well as cell migration, motility and locomotion functions were downregulated (Supplementary Table 6). This indicates that the *Mgp+* fibroblasts are more similar between the adipose tissues in comparison with the *Pi16+* fibroblasts suggesting divergent activities between PVAT and eWAT.

### Spatial transcriptomics mapping of mouse model gene signatures onto human arterial tissues

To examine expression of top mouse DE genes in human tissues, we carried out a spatial transcriptomics experiment using 10X Genomics Visium platform (Fig 3A). Data generated from human aorta and carotid artery samples acquired from three different endarterectomy patients were analyzed with the R-package Seurat. After basic quality control, we mapped the key genes of interest on the human samples (Fig 3B and C). Some genes were highly expressed both in human aorta and carotid arteries (e.g. *CD74*, *APOE*, *MGP*, and *HLA-DRB1*). On the other hand, *JCHAIN* was strongly expressed in aorta but much less in carotid arteries and had an inverse spatial relationship with *APOE* expression, but evident spatial co-expression with several immunoglobulin genes (Fig 3D and E), suggesting plasma cell activity in those regions. PI16 expression was expectedly negligible in all samples, as they strictly contained the aortic or carotid artery wall and no adventitia or other surrounding tissues.

**Figure 3.**
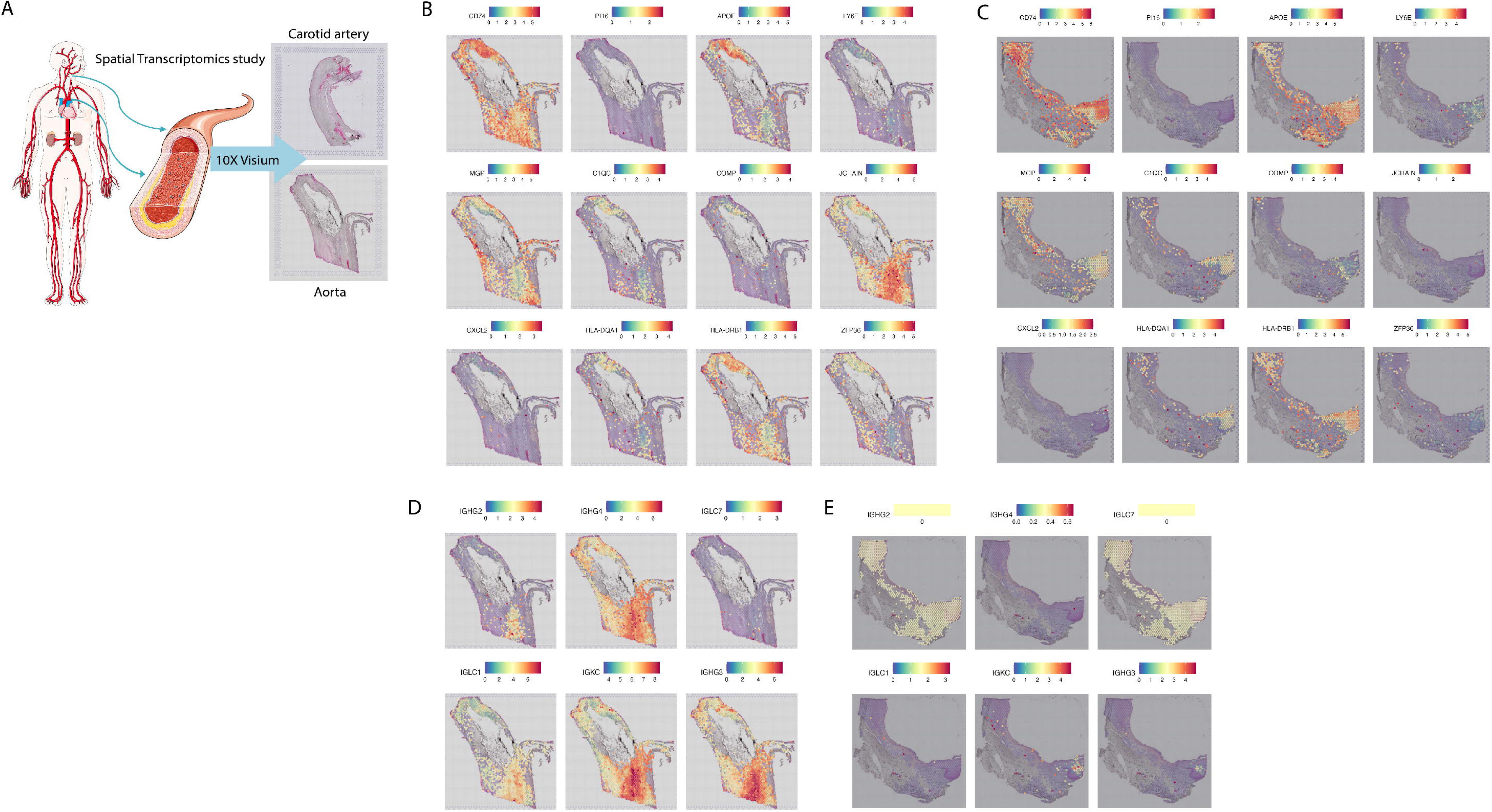
Expression of selected DE genes from the mouse model on representative human arterial tissues (genes: *CD74, PI16, APOE, LY6E, MGP, C1QC, COMP, JCHAIN, CXCL2, HLA-DQA1, HLA-DRB1, ZFP36*). A) Study design. B) Genes of interest (from the single-cell study) in human aorta, C) genes of interest in human carotid artery, D) Immunoglobulin expression in aorta, E) immunoglobulin expression in carotid artery. The colour bar shows normalized expression, where blue refers to no expression and red the maximum of the normalized expression, which varies between genes and samples. The active area on the Visium slides is 6.5×6.5 mm and each spot is 55 µm in diameter.

### PI16+ fibroblasts are found in aortic adventitia, plaques and within the PVAT, and their numbers are reduced in obese state

To confirm the localization of the *Pi16+* progenitor fibroblasts in target tissues, we used IHC of PI16 on sections from formalin-fixed paraffin-embedded aorta-PVAT samples from ten individual mice. The PI16 expression was clearly localized in the fibroblasts in specific patterns around the adipose tissue in the PVAT and the adventitia of the aorta as expected from a previous study^13^ (Fig 4A and B). Although the patterning was observed in both disease model states, there were differences between the non-obese and obese mice.

**Figure 4.**
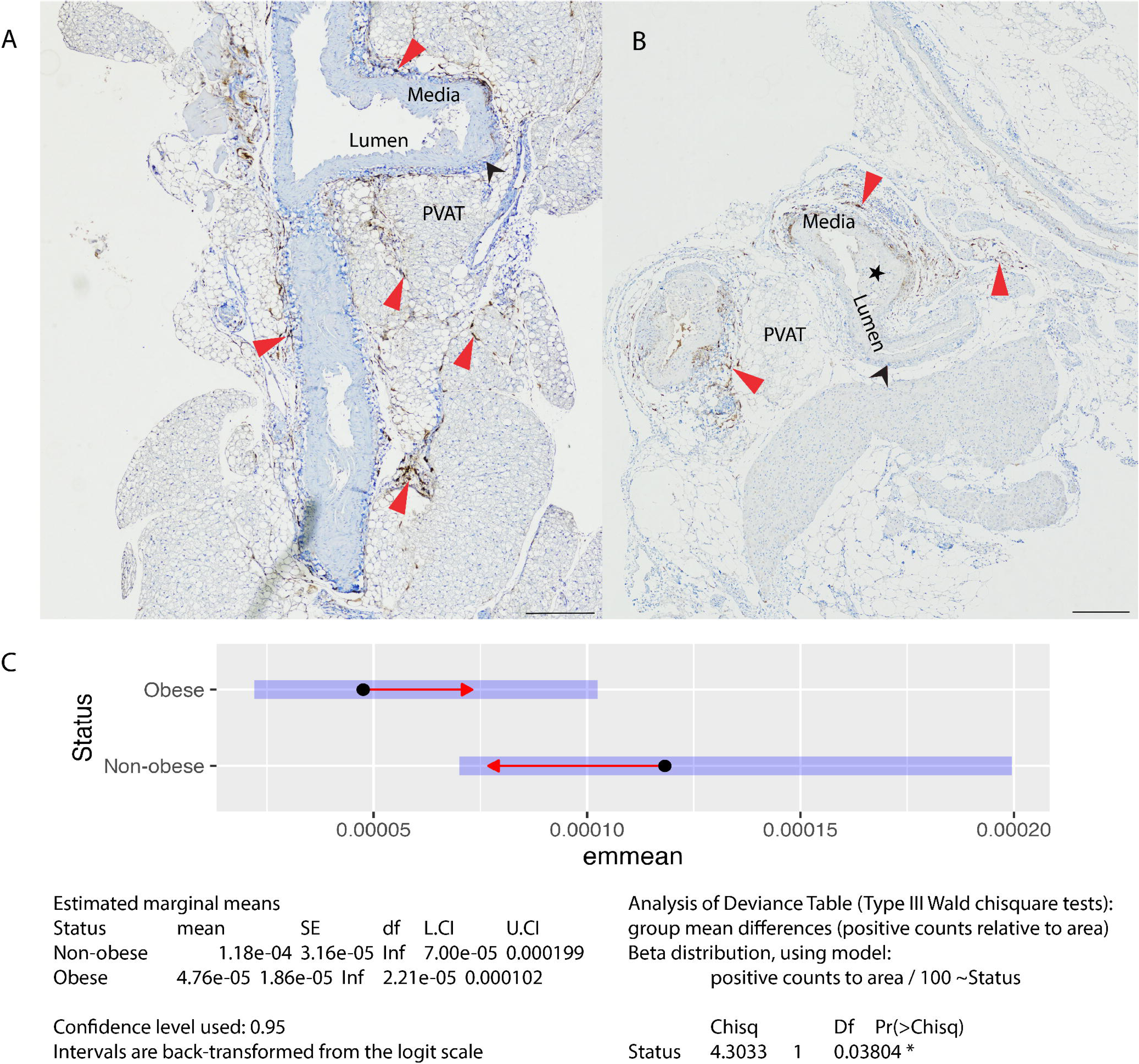
PI16 IHC staining on non-obese (A) and obese (B) mouse PVAT/aorta tissues. Tunica media, lumen, and PVAT are marked with text, adventitia is marked with a black arrowhead, representative PI16+ fibroblasts are marked with red arrowheads, and plaque is marked with a black star. Both scale bars are 250 µm. C) Group mean differences of PI16+ cell counts relative to the area of tissue. Statistical test was Type III Wald chi square test. P-value = 0.03804 (N= 5 obese and 5 non-obese mice). Confidence intervals shown as L.CI (lower) and U.CI (upper). Red arrows represent comparison between the confidence intervals, demonstrating that the difference is significant (arrows are not overlapping each other).

We had seen diminished numbers of *Pi16+* fibroblasts in the obese PVAT-derived single-cell sample, but we wanted to validate this observation at the tissue level. This was done using the same IHC staining images of PI16 as mentioned above. Due to differences in section sizes, we calculated the number of positive cells in relation to the area (mm^2^). To investigate the group differences, we fitted a generalized linear model (postoarea /100 ∼ Status) with beta distribution, and then compared the estimates using a type III Anova, and finally used estimated marginal means to examine how the estimates differ from each other. The estimated group means were significantly different (P-value = 0.03804) (Fig 4C).

### Trajectory analysis confirms Pi16+ fibroblasts as progenitors in both adipose tissues

We performed a trajectory analysis to confirm the progenitor status^14^ of the *Pi16+* fibroblasts and to investigate their developmental relationship with the other adipose tissue-derived fibroblast subpopulations. The analysis was carried out separately for PVAT and eWAT, and the original clustering was used as a priori in the model. The pseudotime trajectory inference corroborated our hypothesis of the *Pi16+* fibroblasts’ progenitor status in both tissues (Supplementary Fig 4A and B). The cell connectivity in both tissues had a similar pattern, where the highest connectivity was around the cluster of *Ccl11+* and *Mfap4+* fibroblasts, indicating that they represent key transition state(s) in the model, while the *Pi16+* progenitors, and the *Mgp+* and *Il1b+* fibroblasts were the start and end points in the trajectory, respectively (Supplementary Fig 4A and B).

### Regulatory network inference analysis reveals tissue and cell-type specific regulons in adipose tissue-derived fibroblasts

Selective activation of different regulons (i.e. transcription factors and their co-regulated target genes) is considered as a major factor in determining differences in cellular activity in different tissues and disease states in complex disorders. Therefore, we carried out a regulatory network inference analysis with pySCENIC to map the transcription factors (TF) that may drive the obesity-associated states in fibroblasts and therefore impact the disease progression in the different tissues.

We found tissue and obesity state specific regulons in the different fibroblast populations, the most interesting being the *Ccl11+* and *Pi16+* subpopulations, having both strong tissue and obesity-state specific regulons, while for instance, *Mgp+* fibroblasts had much less pronounced changes in their TF activity (Fig 5A). The strongest regulons in the full data, based on the area under recovery curve (AUC), are depicted in Figure 8.

**Figure 5.**
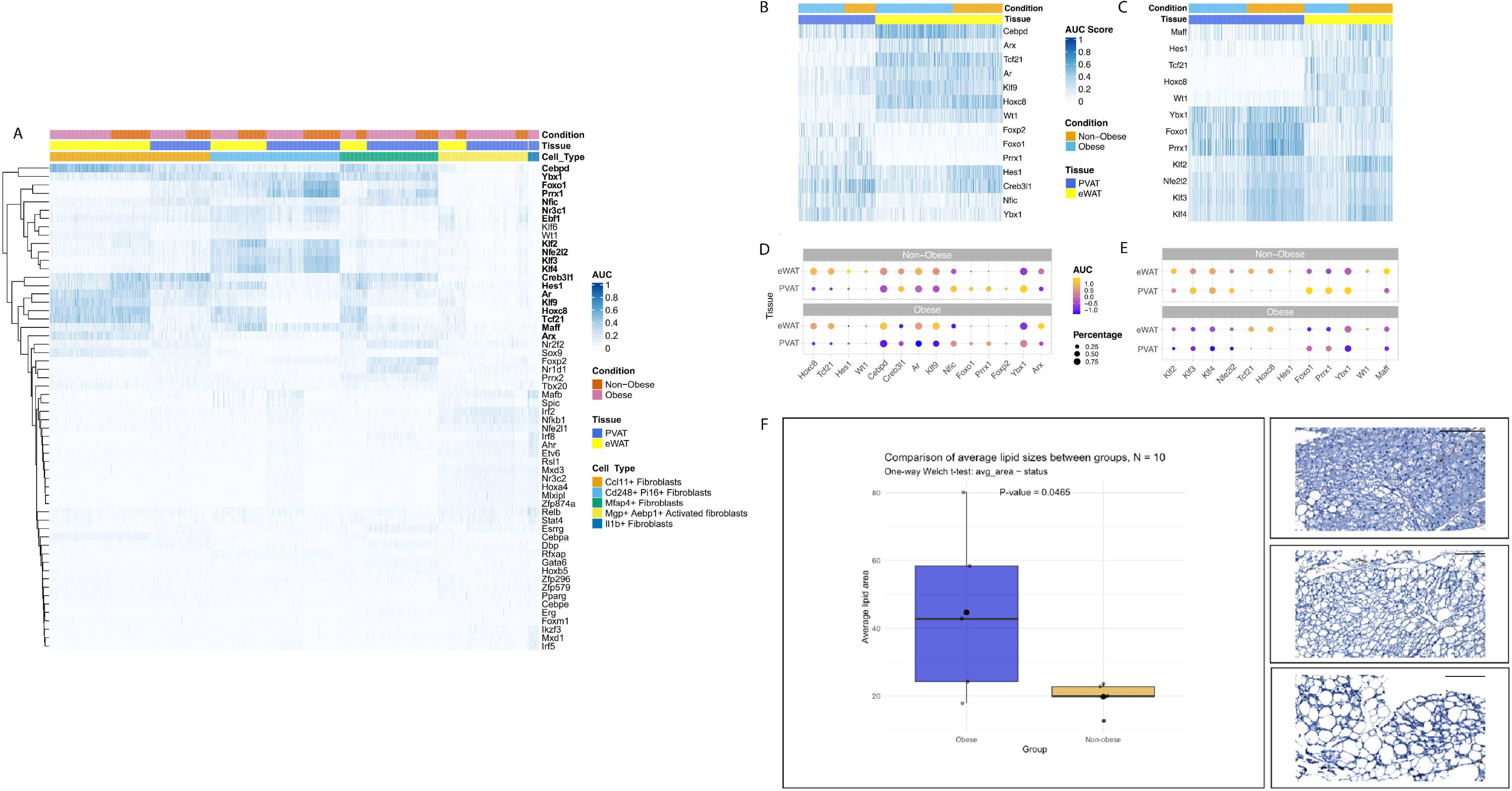
Results of the regulon activity and morphometric analysis. A) Heatmap of the AUC with the top regulons in all of the tissues and fibroblast subsets. The highlighted (bolded) genes are CCAAT enhancer binding protein delta (*Cebpd*), Y-box binding protein 1 (*Ybx1*), Aristaless related homeobox (*Arx*), MAF bZIP transcription factor F (*Maff*), Androgen receptor (*Ar*), Kruppel like factor 9 (*Klf9*), Homeobox C8 (*Hoxc8*), Transcription factor 21 (*Tcf21*), CAMP responsive element binding protein 3 like 1 (*Creb3l1*), Hes family BHLH transcription factor 1 (*Hes1*), Forkhead box protein 1 (*Foxo1*), Paired related homeobox 1 (*Prrx1*), Nuclear factor I C (*Nfic*), Nuclear receptor subfamily 3 group C member 1 (*Nr3c1*), EBF transcription factor 1 (*Ebf1*), Kruppel like factor 2 (*Klf2*), NFE2 like bZIP transcription factor 2 (*Nfe2l2*), and Kruppel like factors 3 and 4 (*Klf3* and *Klf4*). B-C) Heatmaps of the AUC with the top regulons in *Ccl11+* fibroblasts and *Pi16+* fibroblasts in eWAT and PVAT with the two obesity states. D-E) Dot plots of the AUC with the top regulons in *Ccl11+* fibroblasts and *Pi16+* fibroblasts in eWAT and PVAT with the two obesity states. Colour of the AUC shows if the TF is up-(yellow) or down-regulated (blue). Size of the dot refers to the percentage of the cells that express this TF. F) Left side: Average lipid size comparison between the obesity groups (N= 5 obese and 5 non-obese mice) in the perivascular adipose tissue. One-way Welch t-test, P-value = 0.0465. The obese group is coloured blue and non-obese golden. Right side: representative changes in PVAT morphology (Top: normal PVAT in a non-obese mouse, middle: PVAT with slightly abnormal WAT-like morphology in an obese mouse, bottom: PVAT with highly abnormal WAT-like morphology in another obese mouse). The scale bars are all 120 µm.

Next, we investigated the *Ccl11+* and *Pi16+* fibroblasts more closely. In the *Ccl11+* fibroblasts *Cebpd*, *Tcf21*, *Hoxc8*, and WT1 transcription factor (*Wt1*) were predominantly active in the eWAT, showing no significant variation between different obesity states (Fig 5B). All these genes are important for sustaining the WAT phenotype, but *Cebpd* is also an important regulator of immune and inflammatory response related genes^15^. Conversely, Forkhead box P2 (*Foxp2*), *Foxo1*, *Prrx1* were the strongest regulons in PVAT and almost non-existent in eWAT. We were also able to capture regulatory differences between obesity states in these fibroblasts (Fig 5B). *Hes1* had the highest AUC score in the non-obese state in eWAT, while *Creb3l1* had high AUC scores in non-obese state in both adipose tissues.

In *Pi16+* fibroblasts the strongest regulons were *Ybx1*, *Foxo1*, *Prrx1*, and *Klf2-4* (Fig 5C). Notably, all of these genes are associated with adipose differentiation, and *Ybx1* specifically with brown adipogenesis^16^. We observed that the first three are clearly more active in the non-obese state in the PVAT (*Klf2-4* also in eWAT), while *Hes1*, *Tcf21*, *Hoxc8* and *Wt1* were almost exclusively activated in the eWAT but without differences between the obesity states.

### Morphometric analysis reveals significant phenotype changes in PVAT

Our SCENIC analysis indicated strong modulation of adipose tissues during obesity. In line with this, we observed remarkably diverging morphology of the PVAT between the obese and non-obese mice, potentially reflecting a pathogenic conversion^17^. In the obese mice, the PVAT showed transformation toward WAT phenotype. We observed a shift from multilocular PVAT to more unilocular morphology, which is typical to WAT, and increased lipid accumulation was evident (Fig 5D right side), although individual differences were also seen in both groups. To quantitatively assess these differences between the groups, we carried out a morphometric analysis. Using one-way Welch t-test to evaluate whether obese mice have a greater average lipid droplet size than the non-obese mice, we observed that the difference was significant (p = 0.0465, Fig 5D, left side).

### PI16+ fibroblasts may interact with multiple stromal and immune cell types in adipose tissues

Previous studies have shown that PI16 fibroblasts actively interact with endothelial cells but also with immune cells such as macrophages^18,19^. Therefore, we investigated if the PI16+ fibroblasts are co-localizing with T cells and macrophages, which would strengthen the hypothesis of their interactions in these tissues and their possible antigen presentation activity in our model. To achieve this, we carried out a multiplexed immunofluorescence (mIF) staining with PI16, CD3, CD4, and F4/80 (Adhesion G Protein-Coupled Receptor E1) antibodies. PI16 expression was localized in the adipose tissue and the adventitia of the aorta, as well as within plaques, resembling the PI16 IHC staining of the same tissues (Fig 4, Supplementary Fig 5). CD3, CD4 and F4/80 positive cells were evident in the lymph nodes, as expected, and IgG isotype control showed minimal background staining (Supplementary Fig 6 and 7). Lastly, we could clearly observe that the PI16+ fibroblasts were in close proximity to both T cells and macrophages in the PVAT and adventitia, suggesting potential interaction between these cells (Fig 6, Supplementary Fig 5).

**Figure 6.**
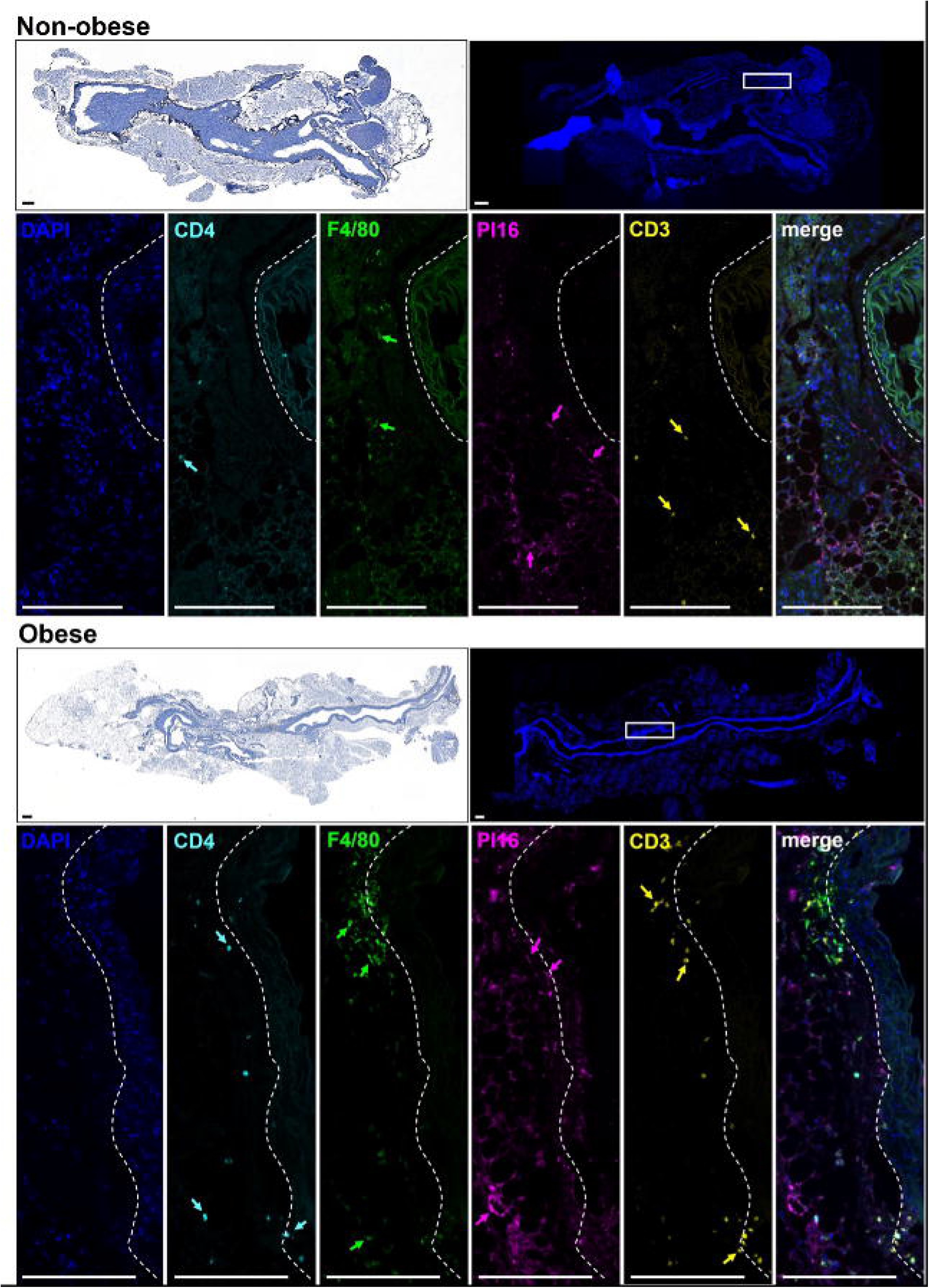
Multiplex immunofluorescence (mIF) staining of PI16 and multiple immune cell markers in PVAT/aorta tissue. Representative images of mIF staining in non-obese and obese mouse tissue showing CD4, F4/80, PI16 and CD3 expression in the PVAT and the adventitia of the aorta (white dashed line). Images of the PI16 IHC staining (upper left panels) depict the morphology of the mIF stained tissues (not adjacent sections). DAPI (4’, 6-diamidino-2-phenylindole) was used as nuclear counterstain. Scale bar 200 µm.

Next, we explored other potential cell-cell interactions involved in the regulation of the PVAT-derived *Pi16+* progenitor fibroblasts. We used a method, which assesses how ligands expressed by sender cell types predict expression changes in the receiver cell type, enabling inspection of the cellular interactions not only between ligands and receptors, but also downstream target genes potentially regulated by the receptors within the receiver cells. The *Pi16+* fibroblasts were defined as the receivers and the sender cell types were the cells that were observed in the adipose tissues and showed significant differential gene expression between the obesity states (see Methods).

The results revealed multiple general ligands and one endothelial cell specific ligand (*Vwf*; Von Willebrand Factor), that could target a set of genes downstream (Supplementary Fig 8A). The most active general ligand was *Apoe*, followed by *Tgfb2* (transforming growth factor beta 2) (Supplementary Fig 8B), and both had multiple predicted target genes with strong regulatory potential (Supplementary Fig 8C). *Apoe* was expressed to some extent in many sender cell types, but had strong average expression in different macrophages, while *Tgfb2* was mainly expressed by a small proportion of other fibroblast populations. *Vwf* was only expressed by endothelial cells and some mesothelial cells (Supplementary Fig 8D). The target genes towards which *Apoe, Tgfb2,* and *Vwf* had the highest regulatory potential are listed in Table 2. Another interesting ligand was *Il1b*. It wasn’t among the most active ligands, but it showed strong regulatory potential (Supplementary Fig 8B and C) with multiple target genes. In this analysis, it was most strongly expressed in a specific granulocyte population, but also by different DC and monocyte populations (Supplementary Fig 8D).

**Table 2.**
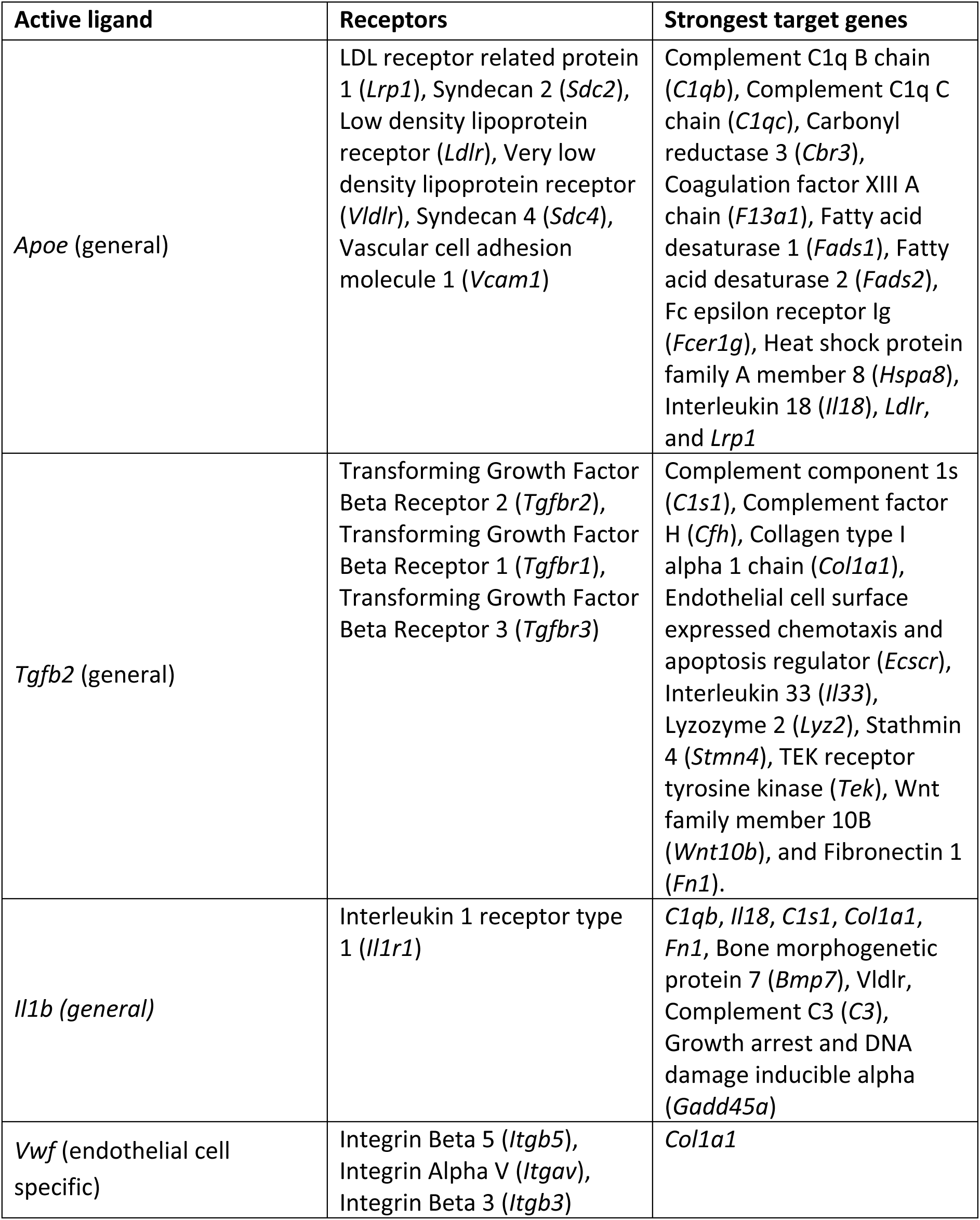
The most active ligands and their receptors and strongest target genes from the cell-cell interaction analysis.

## Discussion

Together, our application of 5ʹ single-cell RNA sequencing and rigorous validation approaches have enabled the identification of key changes within perivascular and epididymal adipose tissues in a murine model of atherosclerosis during obesity and normal weight. Driven by our data, we have focused on different fibroblast subpopulations and especially on *Pi16+* fibroblasts in this study. These fibroblasts have previously been determined as a progenitor fibroblast type^13,18^, giving rise to many other fibroblast populations, including myofibroblasts, and can act as a precursor for adipocyte lineage, thus playing a role in adipogenesis^18^. Our trajectory analysis confirmed this also in this atherosclerosis mouse model. However, as suggested before, *Pi16+* fibroblasts are not solely progenitors to other cell populations but can have multifaceted functions during steady state as well as in inflammation^18^. They have also been shown to promote immune cell trafficking in different contexts ^18,20^, and to interact with specific cytotoxic T cells and macrophages within perivascular niches in cancer^20^.

Our differential gene expression analysis revealed genes that were upregulated in obesity in multiple cell types across aorta, PVAT and eWAT, such as *Cd74* and *Apoe*. Both genes have been previously strongly linked to atherosclerosis^21,22^. In humans, the production of CD74 protein is elevated in carotid atherosclerotic plaque shoulder regions, with high macrophage accumulation, and the production of CD74 is also upregulated in vascular smooth muscle cells and monocytes under interferon gamma stimulation ^23^. Indeed, our spatial transcriptomics data showed that the strongest *CD74* expression was in regions with accumulation of the macrophage marker *CD68* (Supplementary Fig 9). Similar CD74 expression patterns appeared in our mouse PVAT-aorta samples using IHC, confirming its expression in fibroblasts (Supplementary Fig 10). *CD74+* cancer associated fibroblasts function as antigen presenting cells in multiple cancer types^20^, while cardiac fibroblasts participate in antigen presentation and contribute to cardiac fibrosis in mouse models^24^. Our gene set enrichment analysis indicated that PVAT-derived *Pi16+* and *Mgp+* fibroblasts had antigen presentation and processing functions during obesity. Therefore, we suggest that these fibroblast populations in PVAT might act as antigen presenting cells via MHC II and may participate in T cell mediated immune responses in atherogenic environment during obesity. Further functional studies are needed to demonstrate this *in vitro*.

Even though the role of APOE in atherosclerosis is undeniable, APOE expressing fibroblasts remain uncharacterized in atherosclerosis or obesity. To our knowledge, this is the first time *Apoe* expression has been detected in adipose tissue-derived fibroblasts in an atherosclerosis model, and its function in these cell populations warrants further research. The other cell types that had significantly increased *Apoe* expression in the obese state were an aorta-derived *Fscn1+ Apol7c+* dendritic cell population (Fascin 1 has previously been linked to migrating and mature DCs^25^) and PVAT-derived intermediate monocytes. Apoe deficiency has been linked with increased antigen presentation in DCs and enhanced T cell activation^26^. Apoe is important for normal antigen presentation and processing functions of DCs via membrane lipid homeostasis; MHC II molecules concentrate in lipid rafts, which will accumulate abnormally if DCs do not express Apoe^26^. We suggest that Apoe might have a similar role in antigen-presenting fibroblasts in building lipid rafts, in which the MHC II molecules then concentrate, which could explain its upregulation in the PVAT-and eWAT-derived fibroblasts that demonstrated antigen presentation/processing potential. Moreover, in monocytes, where classical and intermediate monocytes are the main producers of APOE^27^, its overexpression can lead to decreased antigen presentation capacity by monocytes^28^, highlighting APOE’s importance to functional antigen presentation.

Interestingly, multiple complement cascade component genes were also upregulated in different cell types and tissues in the obese state of our mouse model. Most prominently *C4b* was upregulated in eWAT-derived *Mgp+* fibroblasts, and *C1qb* and *C1qa* in PVAT-derived *Pi16+* fibroblasts. The impact of the complement system in atherosclerosis is well-studied and has both atherogenic and-protective effects, depending on the stage of complement activation^29^. However, knowledge of local cell-specific expression is lacking, and it may be the key to understanding how complement system drives atherosclerosis^29^. Previously, a single-cell RNA sequencing study of human atherosclerotic plaques indicated one *CD68+* myeloid cell population that had high expression of *C1QA*, *C1QB* and *C1QC*^30^. Another study showed that *C3* expression was high in VSMCs from a proximally adjacent region of a plaque from carotid artery samples^31^. We are the first to show that complement component expression might also happen in fibroblasts in the context of atherosclerosis during obesity, although this requires further validation. Moreover, we observed multiple complement cascade components expressed in the human aorta and carotid arteries with distinct spatial patterns (Supplementary Fig 11).

5’ RNA sequencing enables the interrogation of the transcription start sites and therefore the study of transcription factor expression profiles. We found differences in multiple interesting transcription factors between the obesity states. This analysis also indicated divergent regulation programs of *Ccl11+* and *Pi16+* fibroblasts in the two adipose tissues. The strongest differences between the adipose tissue types were observed for *Cebpd*, *Tcf21*, *Hoxc8*, *Hes1,* and *Wt1*, expressed in the eWAT-derived fibroblasts, while minimally expressed in PVAT. This difference was pronounced in the *Pi16+* fibroblasts. *Cebpd* is an important regulator of genes that participate in immune and inflammatory responses^15^, and it has a central role in early adipogenesis^32^*. Tcf21* is a WAT marker and contributes to the proinflammatory environment and ECM remodelling in visceral adipose tissue^33^. *Ho*xc*8* is another WAT gene, and its suppression leads to browning of WAT^34^. *Hes1* and *Wt1* also regulate WAT formation. *Wt1* is expressed by visceral WAT, whereas subcutaneous WAT and brown adipose tissue (BAT) do not express *Wt1*^35^. Browning of eWAT is observed in *Wt1* heterozygous mice^35^. *Hes1*, which was observed to be a strong regulon in the non-obese eWAT derived *Ccl11+* fibroblasts, is a downstream target of Notch signalling pathway and prevents browning of adipose tissue by modulating expression of *Prdm16* and *Ppargc1a*, which are important regulators of BAT thermogenesis^36^. Finally, *Creb3l1* has been indicated as an inducer of adipogenesis in a study where *CREB3L1* knockdown in human adipose-derived stem cells increased different measures of adipogenesis^37^.

Two key TFs in PVAT that showed higher activity in non-obese state, *Foxo1* and *Prrx1*, regulate cell proliferation and differentiation, and both are linked to mediating myofibroblast differentiation^38,39^. Moreover, *Prrx1* has been shown to be a cell stemness regulator ^40^ as well as to participate in adipocyte differentiation^41^. Knocking down *PRRX1* in human adipose-derived stem cells increase lipid accumulation and may repress adipogenesis^37^. The Kruppel-like factors have been linked with a multitude of cellular processes but most interestingly they have been demonstrated to be strongly involved in adipogenesis and lipogenesis^42^. For example, myeloid-derived Klf2 is a key regulator of obesity and is downregulated during high-fat diets^43^. We also established that its expression was lower in both adipose tissues of the obese state mice across all the fibroblasts, and specifically in the *Pi16+* fibroblasts. Ultimately, we were able to detect the putative adipose tissue modulators, and to demonstrate that the transcriptional regulation of *Pi16+* fibroblasts had changed notably between the obesity states. This was supported by the morphometric analysis (Fig 5D), which showed that the morphology of the PVAT had significantly changed. These results indicate that obesity has induced a remarkable shift in the PVAT functionality and has driven the progenitor as well as other fibroblasts toward immunoregulatory phenotypes, which may actively participate in immune responses in the perivascular environment.

We also performed a cell-cell interaction analysis to study how and with what cells the PVAT-derived *Pi16+* progenitor fibroblasts might interact. *Apoe* was again highlighted, its target genes being, for instance, C1 complement cascade members, some essential fatty acid desaturases, low-density lipoprotein receptor proteins, and interleukin 18, a proinflammatory cytokine (Table 2). Complement cascade members are clearly important, and the fatty acid desaturases have some interesting immune-related functions. For example, *Fads2* is essential for proper leukotriene production during immune responses in macrophages^44^. Lipoprotein receptor signalling, represented here by *Ldlr* and *Lrp1*, is one of the focal pathways affecting atherosclerosis development^45^, while *Il18* is linked to atherosclerosis plaque instability^46^, and interferon gamma-dependent acceleration of atherosclerosis^47^. *Il18* was also one of the target genes of *Il1b* in this analysis. Another interesting target for *Il1b* was *Bmp7*, which is abundant in lipid-rich plaques and intensifies monocyte extravasation in humans^48^, a central process in atherogenesis. Generally, *Tgfb2* and *Il1b* had multifaceted targets of both ECM component and immune related genes (Table 2). For instance, *Il33*, one of the targets of *Tgfb2* is atheroprotective and it has been suggested that this protective process could be due to inhibition of cell adhesion^49^, possibly decreasing the adhesion and migration of immune cells into the arterial wall.

The endothelial cell-specific ligand *Vwf* was especially intriguing, because *Pi16+* fibroblasts have been functionally shown to interact with endothelial cells to promote transendothelial leukocyte trafficking^18^, which could possibly be explained by the pivotal role of *Vwf* to endothelial permeability in vasculature^50^. It remains to be shown *in vitro*, how *Pi16+* fibroblasts and endothelial cells may orchestrate leukocyte trafficking in atherosclerosis.

Altogether the interaction analysis revealed that PVAT-derived *Pi16+* fibroblasts may impact atherosclerosis through multifaceted roles in obesity.

Our study had some limitations. First, only male mice were used in this study. In the future it would be important to extend similar studies to include both sexes to define how atherosclerosis may be differentially driven in them. Second, the lack of human adipose tissue samples; our spatial transcriptomics data included only aorta and carotid arteries without adjoining tissues, with limited number of human patients. Still, we showed how some of the most interesting genes from the single-cell study were expressed in the human tissues. And finally, the composition of PVAT varies by region and our validation sections used for IHC and mIF mostly represented the abdominal part of the descending aorta. Future studies should investigate if there are significant site-specific differences. Nonetheless, our single-cell data was constructed from the aorta and PVAT covering the whole aorta from the ascending aorta to the abdominal aortic region where the renal arteries diverge. Future research should explore how *Pi16+* fibroblasts functionally interact with immune cells and other stromal cells, such as endothelial cells, and how they may orchestrate atherosclerosis progression or inhibition together from perivascular adipose tissues in human patients.

Altogether, we present a comprehensive single-cell and multi-tissue perspective of the immune-stromal landscape in a murine model of atherosclerosis during obese and non-obese states. The combination of 5’ single-cell RNA sequencing and rigorous validation methods defined perivascular adipose tissue-derived fibroblasts as active participants in immune regulation within the tissue, and as potential contributors to atherosclerosis through local stromal-immune interactions. Our study offers insights into how obesity may exacerbate atherosclerosis via extensive changes in the PVAT functionality through different fibroblast populations, but especially by modulating the *Pi16+* fibroblasts.

## Sources of funding

Research Council of Finland (314557, 335977, 335975, 352968), InFLAMES Research Flagship Centre (337530).

## Acknowledgements

Most of the computational work was carried out at the CSC - IT Center for Science (Finland) high performance computing system (https://research.csc.fi/service/puhti/).

We wish to thank the following core facilities of Turku Bioscience Centre (University of Turku, Finland), Univeristy of Eastern Finland (UEF), and Biocenter Finland (https://biocenter.fi/) for their invaluable services and knowledgebase that enabled the different experiments carried out in this study: Histology core facility of the Institute of Biomedicine (histology and antibody staining), Single-Cell Omics core (mIF staining), Cell Imaging and Cytometry core and Medisiina Imaging Centre (imaging, and morphometric analysis), PET-centre and Lab Animal Centre of the University of Eastern Finland (animal work), Finnish Functional Genomics Centre (sequencing services), Medical Bioinformatics Centre in Turku and Single Cell Genomics Core of University of Eastern Finland (bioinformatics consultancy).

We also extend our thanks to Dr. Milla Salonen who provided valuable insight regarding the statistical tests used in the morphometric analysis and group comparisons of PI16 activity in PVAT, and to MSc Sara Metso who helped in imaging of the IHC validation sections.

## Disclosures

The authors declare that they have no conflict of interest.

## Supplemental Material

Figures S1-S13

Supplementary Methods

Tables S1-S6

## Notes

### Competing Interest Statement

The authors have declared no competing interest.

### Summary of Updates

The manuscript has been revised according to reviewer comments.

