## Supplementary Figures S1-S13 and Supplementary Methods for "Immune interaction signatures in adipose tissue fibroblasts in obesity-associated atherosclerosis"

**Supplementary materials**

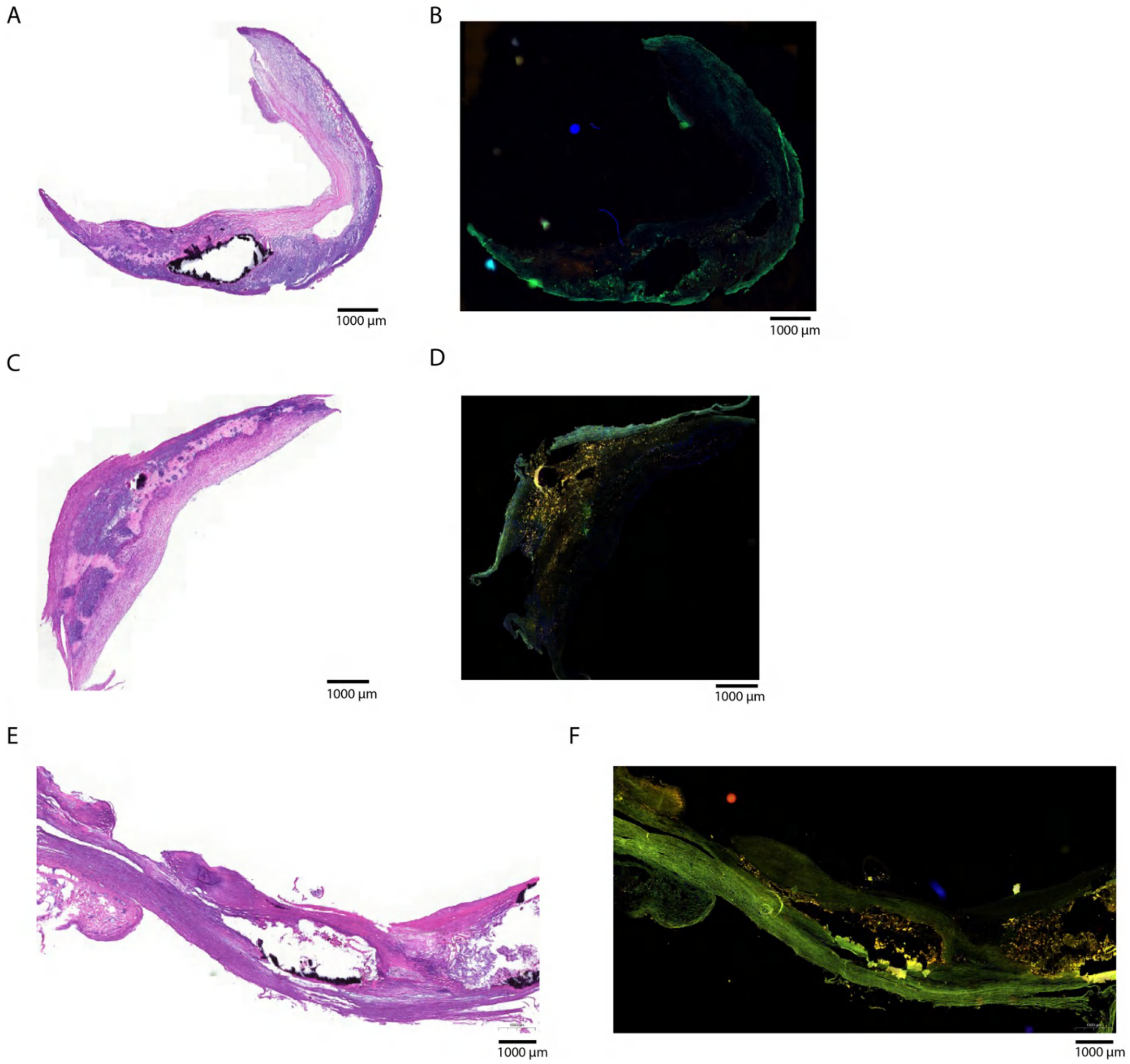


Supplementary Figure 1. CD68, CD206 and H&E staining in human aorta and carotid artery samples used for selecting the regions of interest for spatial transcriptomics. A) H&E staining of carotid artery (patient 1), B) CD68/CD206 staining of carotid artery (patient 1), C) H&E staining of carotid artery (patient 2), D) CD68/CD206 staining of carotid artery (patient 2), E) H&E staining of aorta, F) CD68/CD206 staining of aorta. The scale bars are all 1000 µm.

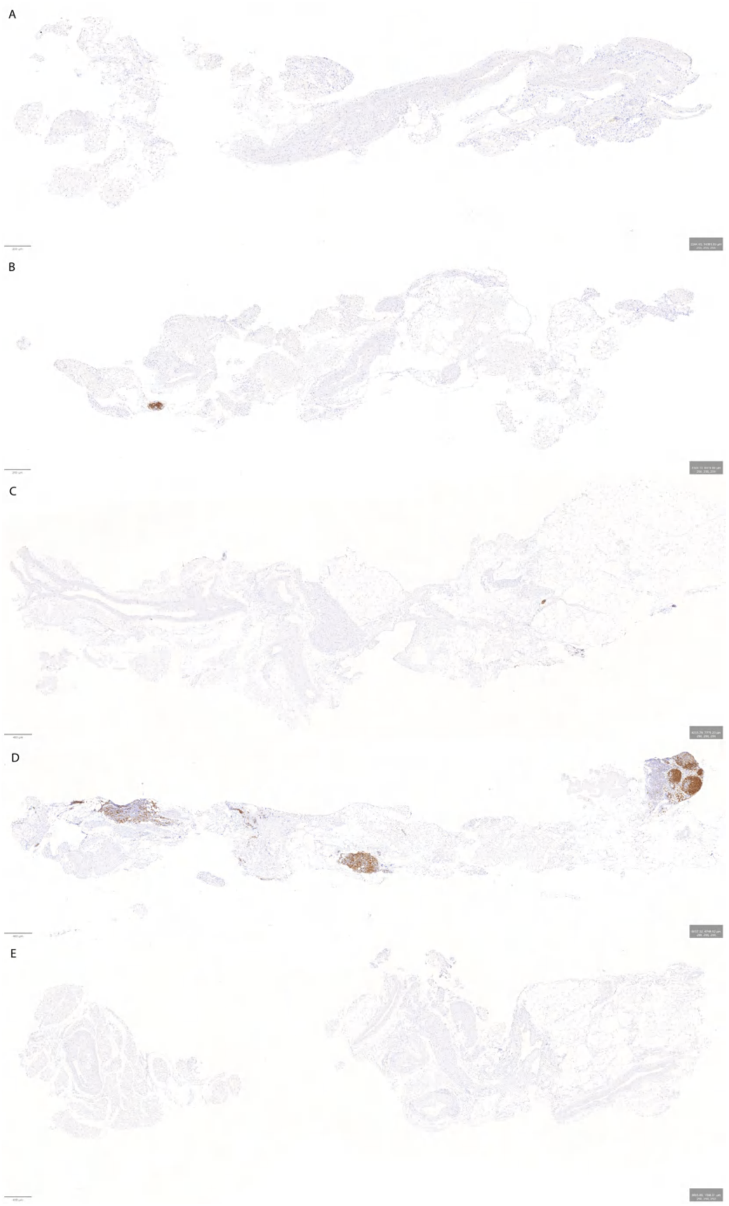

Supplementary Figure 2. Representative IHC staining of CD19 in mouse PVAT-aorta tissues. The protein expression localized to lymph nodes. Sections A-B) are from two non-obese mice (scale bars A=200 µm, B=250 µm), and sections C-E) are from three obese mice (scale bar 400 µm in all). Positive staining (brown) clearly localizes to lymph nodes and - vessels. Full imaging data (N=5+5) is available from 10.6084/m9.figshare.29279045.

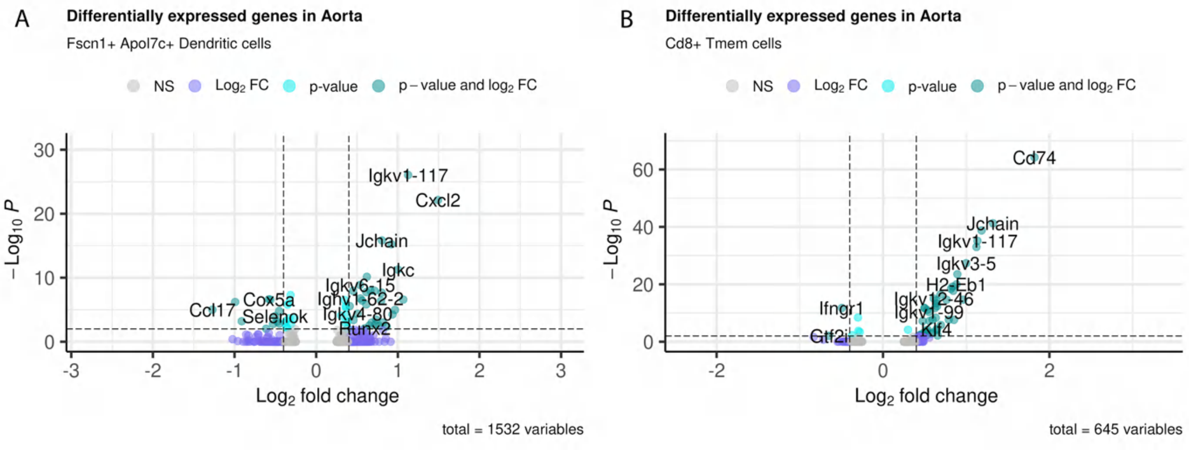


Supplementary Figure 3. Differentially expressed genes in aortic *Fscn1+ Apol7c+* DC (A) and *Cd8+* Tmem cell (B) populations. Positive fold change indicates higher expression in the obese model state. Statistical test was Wilcoxon rank sum test. Threshold for log2 fold change was 0.5 and threshold for significance was adjusted P-value < 0.05.

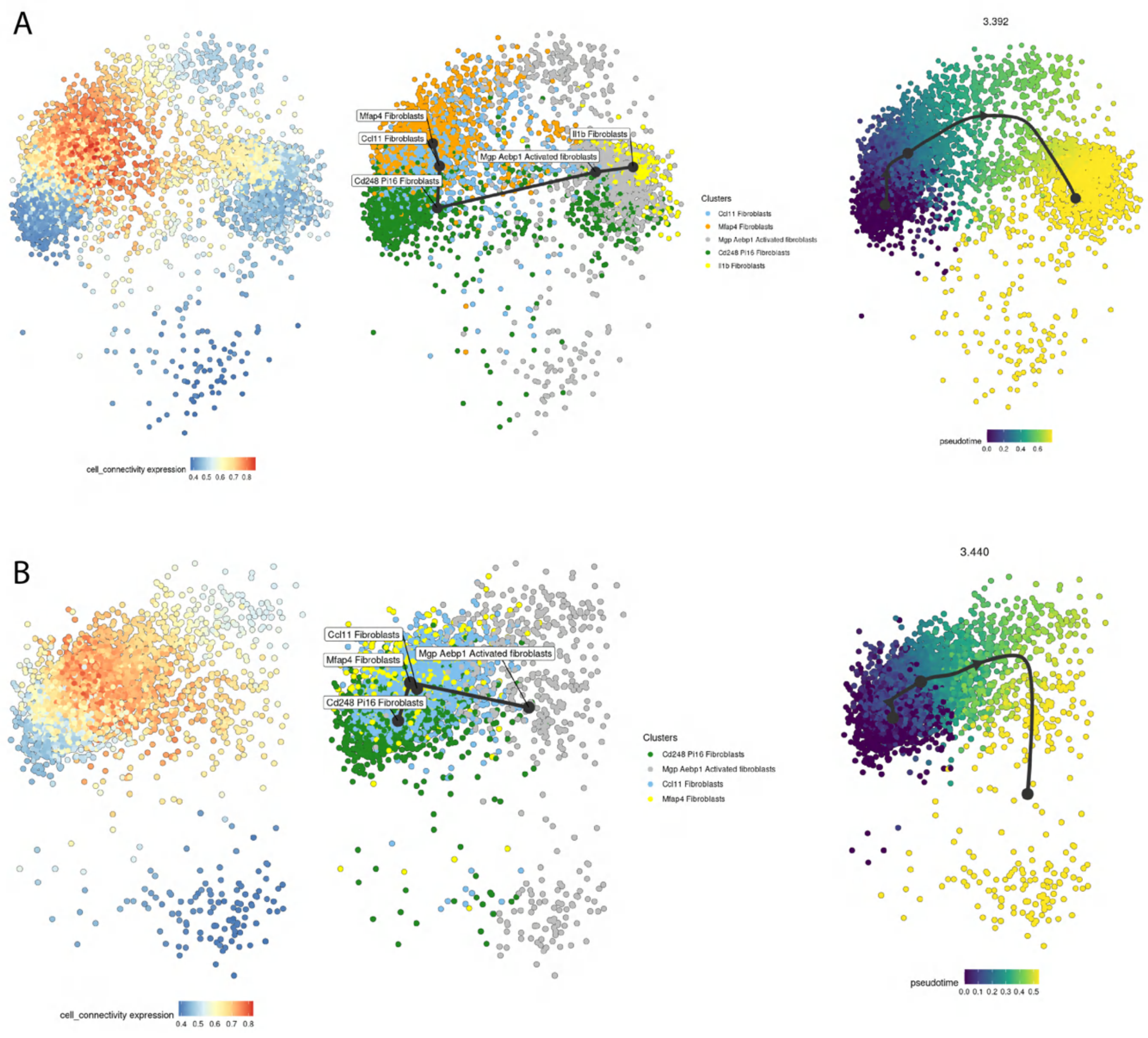

Supplementary Figure 4. Cell connectivity, original clustering, and pseudotime trajectory from the trajectory analysis using Totem R-package. A) PVAT (number of cells = 4740), B) eWAT (number of cells = 3621). In the connectivity plot, the orange and red colours indicate high connectivity and conversely blue low connectivity. The high connectivity cells represent key transition states and and low connectivity refers to start and end points in the trajectory. In the pseudotime plot the dark blue represents the inferred starting point and yellow the end of the pseudotime trajectory.

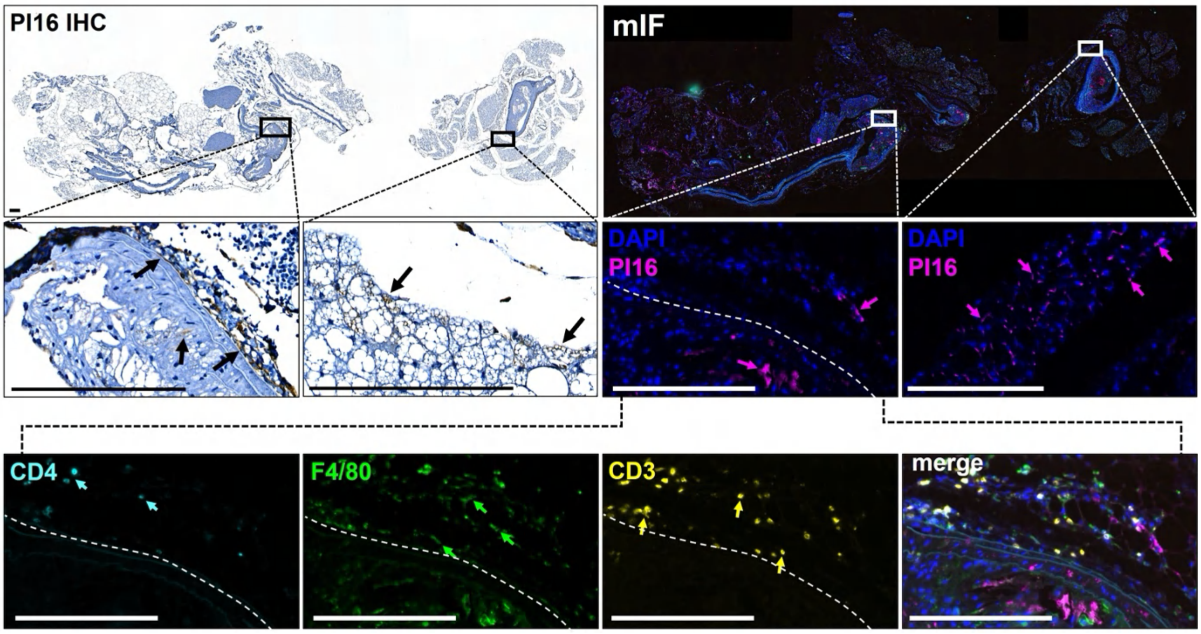


Supplementary Figure 5. Immunohistochemistry (IHC) and multiplex IF (mIF) staining of PI16 and multiple immune cell markers in obese mouse PVAT/aorta. Upper panel: Representative images of PI16 IHC and mIF staining in obese mouse tissue of the same individual (not adjacent sections). Lower panel: CD4, F4/80 and CD3 expression near the plaque and adventitia of the aorta (white dashed line). DAPI (4', 6-diamidino-2-phenylindole) was used as nuclear counterstain. Scale bar 200 µm.

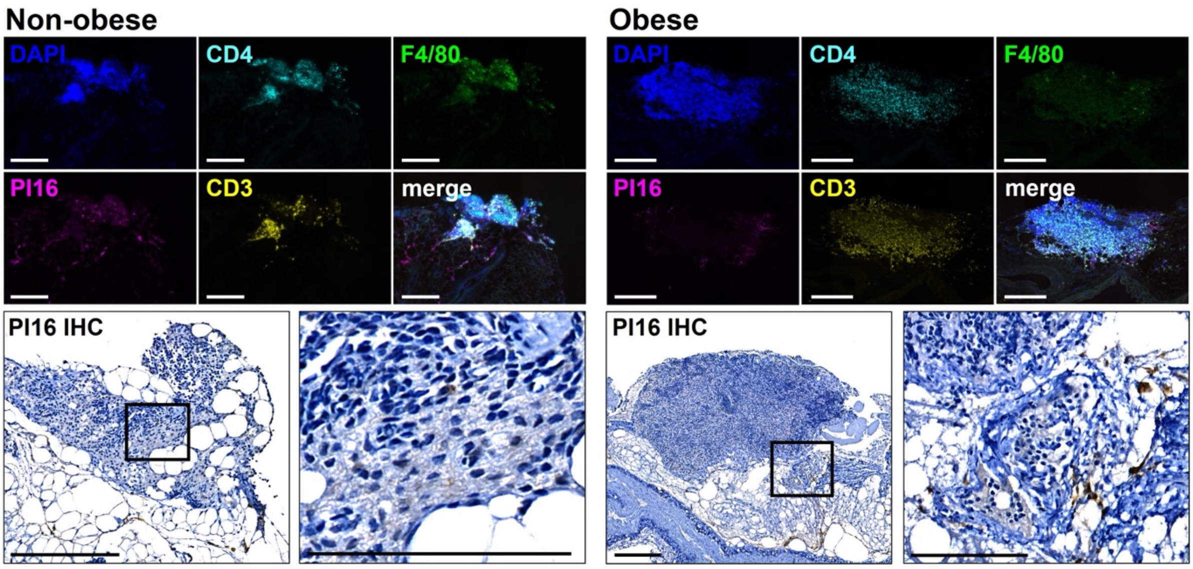


Supplementary Figure 6. Multiplex IF staining of immune cell markers in non-obese and obese mouse lymph nodes. Representative images of CD3, CD4, F4/80 and PI16 expression in mouse lymph nodes by mIF staining. Images of the PI16 IHC staining (lower panels) depict the morphology of the mIF stained tissues (not adjacent sections). DAPI (4’, 6 -diamidino-2-phenylindole) was used as nuclear counterstain. Scale bar 200 µm, and 100 µm in the PI16 IHC close-up.

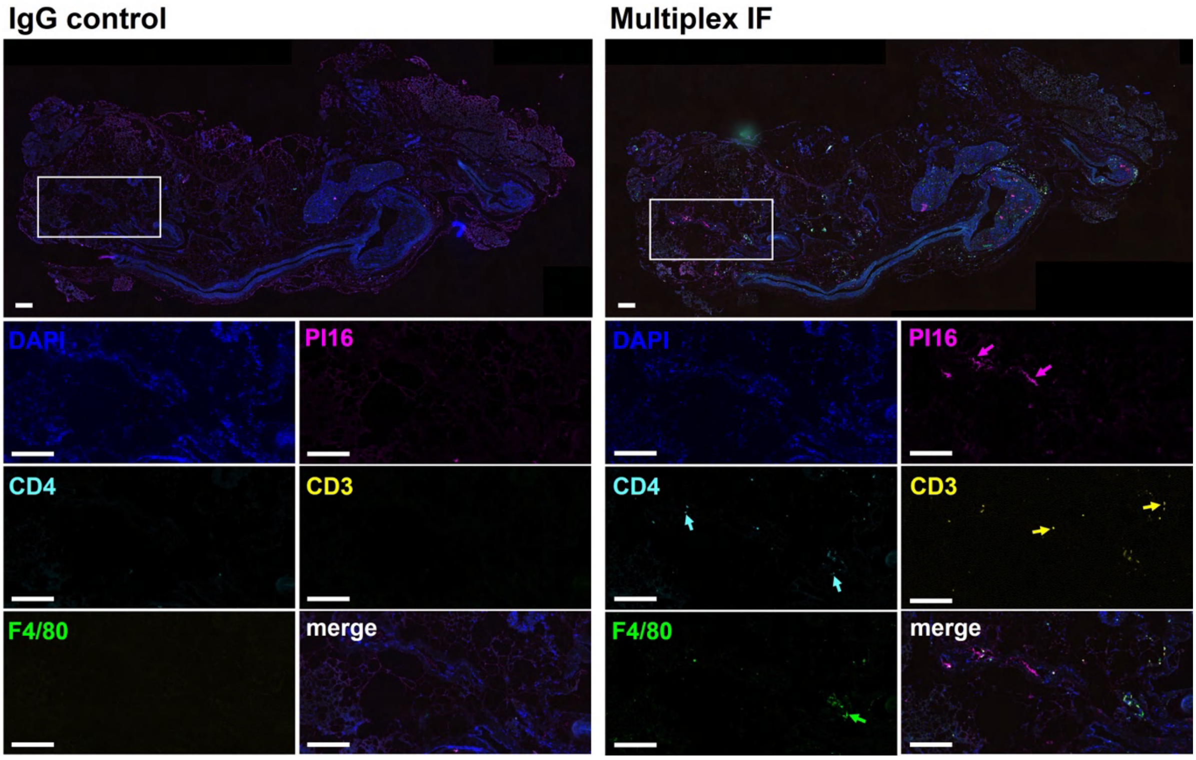


Supplementary Figure 7. Negative control mIF staining of matched adjacent mouse PVAT/aorta tissue. Representative images of PI16, CD4, CD3 and F4/80 mIF staining in obese mouse PVAT/aorta tissue (right panel) and IgG isotype control of adjacent section (left panel). DAPI (4’, 6 -diamidino-2-phenylindole) was used as nuclear counterstain. Scale bar 200 µm.

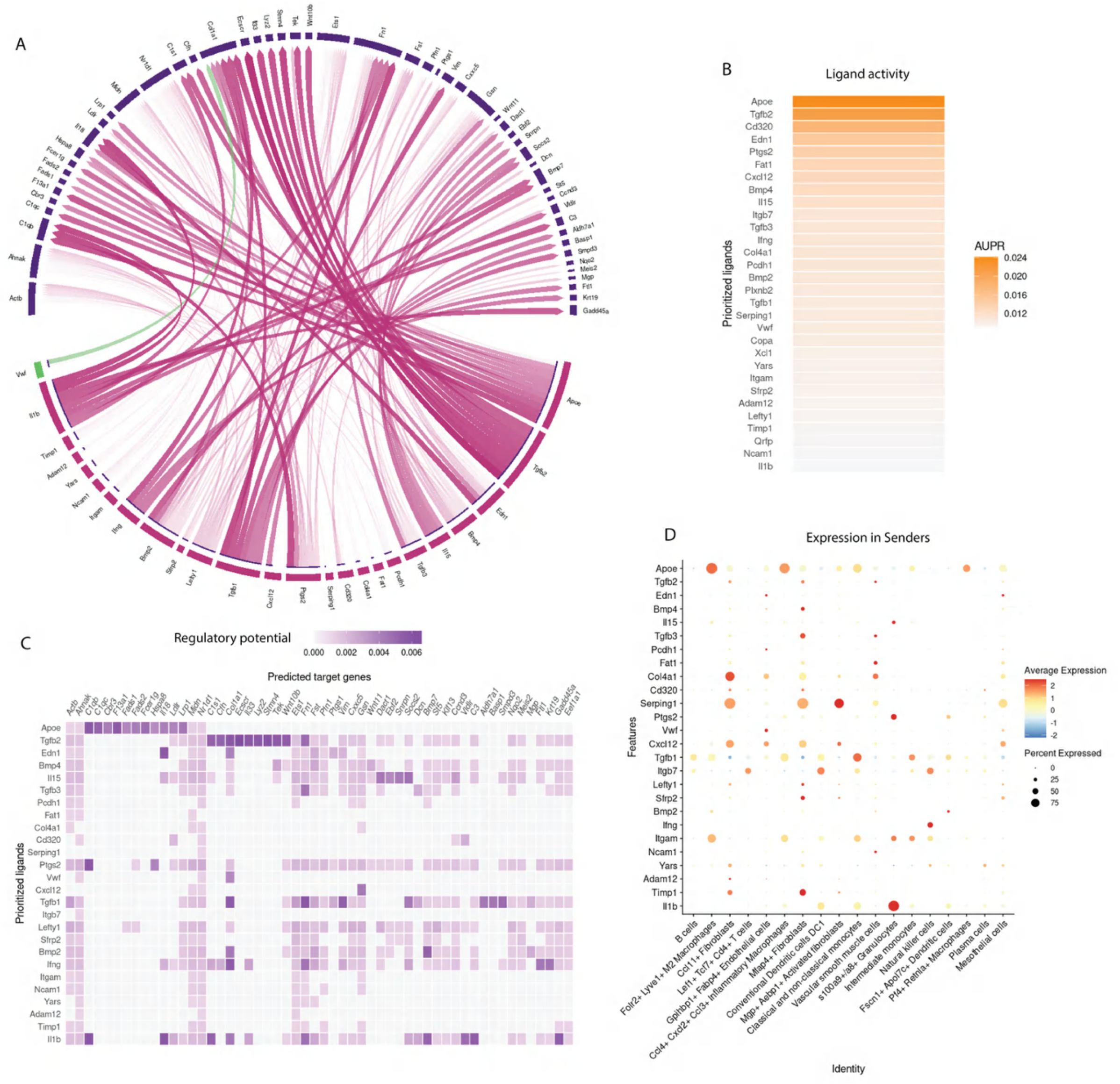


Supplementary Figure 8. Summary of NicheNet cell-cell interaction results. Number of cells in analysis; receiver = 1255, senders =9727. A) Connections between general ligands (burgundy blocks/arrows), one endothelial specific ligand (green block (arrow), and their target genes (dark purple blocks). The stronger the regulatory potential, the stronger the line between the ligands and their targets. B) Ligand activity (AUPR: area under the precision-recall curve). C) Regulatory potential of the predicted ligands against the predicted target genes. D) Ligand gene expression in the sender cells.

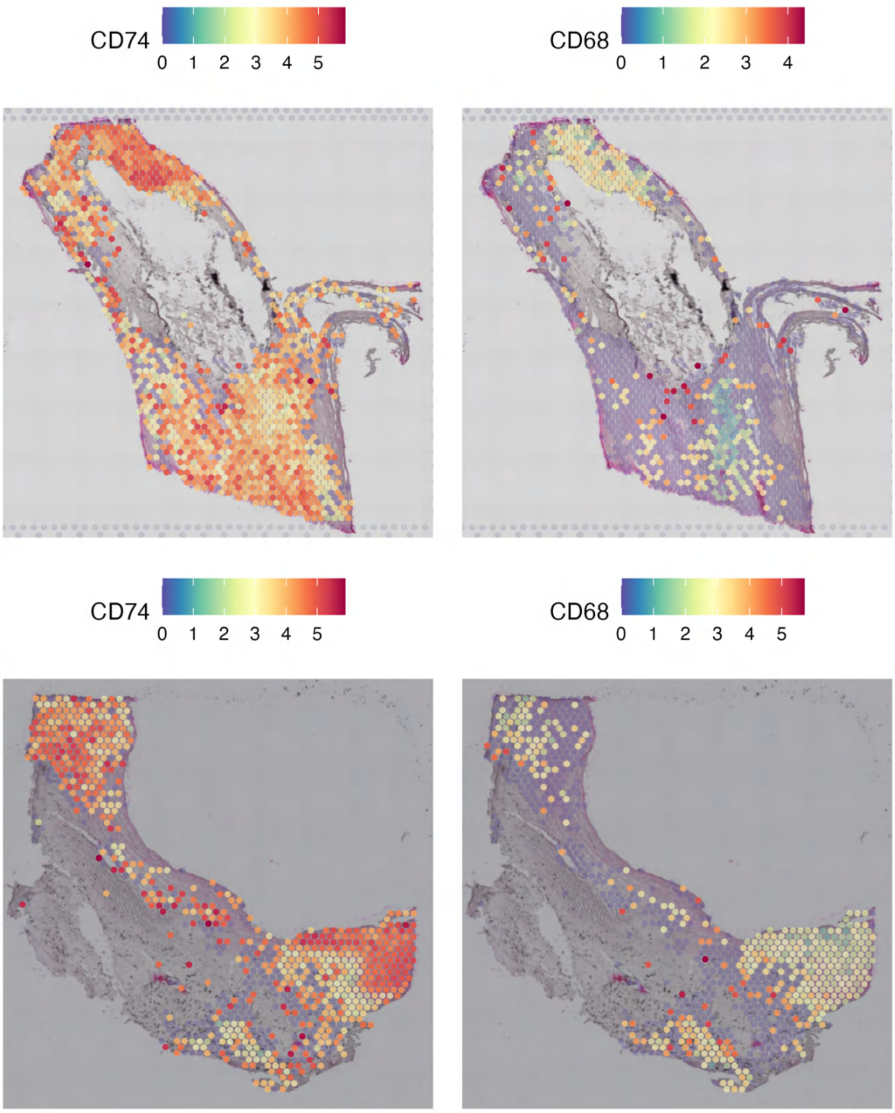


Supplementary Figure 9. Spatial *CD74* and *CD68* expression in human aorta and carotid artery. The colour bar shows normalized expression, where blue refers to no expression and red the maximum of the normalized expression, which varies between genes and samples. The active area on the Visium slides is 6.5x6.5 mm and each spot is 55 µm in diameter.

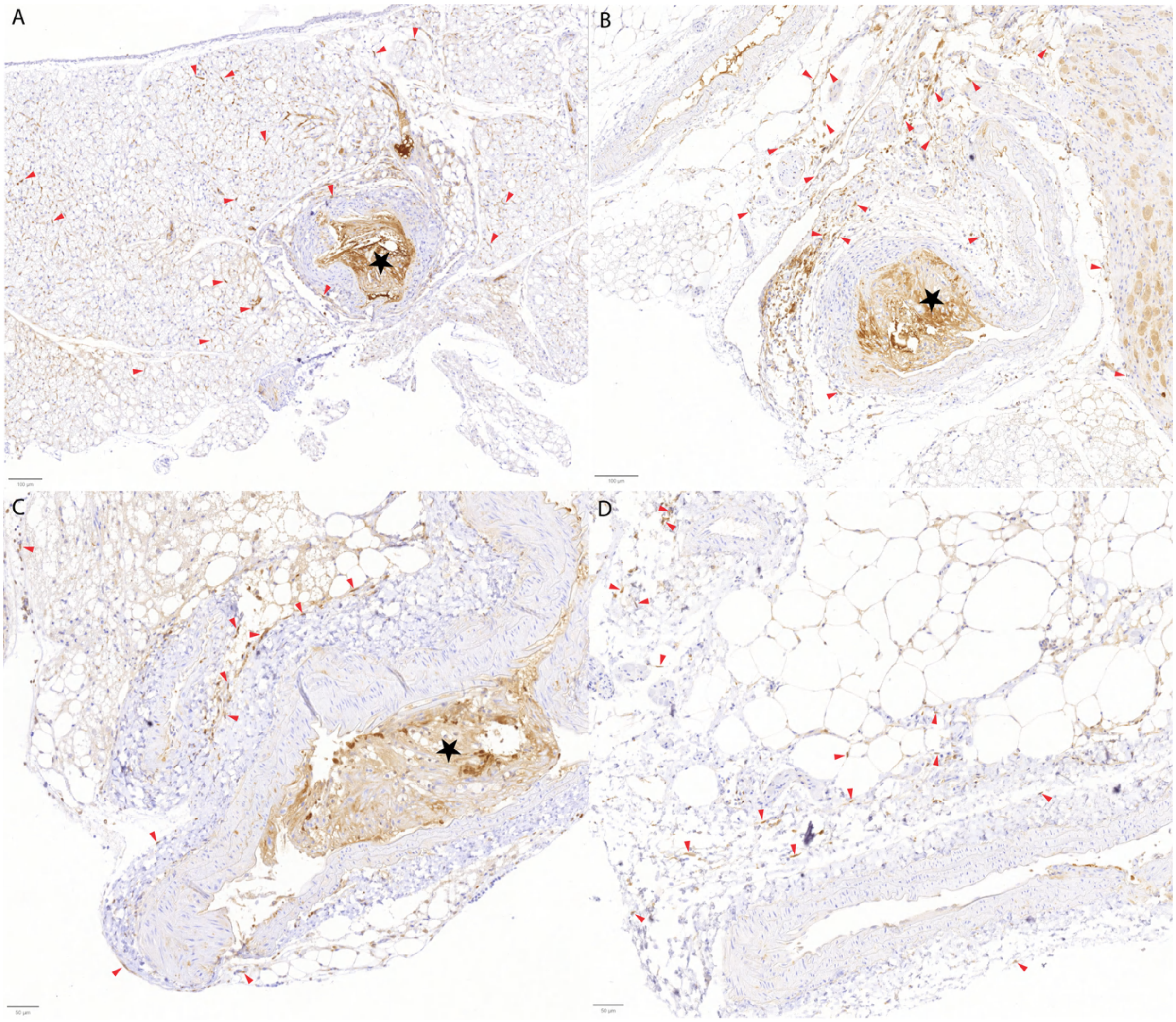


Supplementary Figure 10. Representative images of IHC staining of CD74 in obese mouse PVAT-aorta tissues. The protein expression (brown) localized strongly to plaques (marked with black stars), but we confirmed that CD74 was also expressed by fibroblasts (representative cells marked with red arrowheads). The scalebar is 100 µm for the upper figures and 50 µm for the lower figures. Full imaging data (N=5+5) is available from [10.6084/m9.figshare.30030004](https://doi.org/10.6084/m9.figshare.30030004).

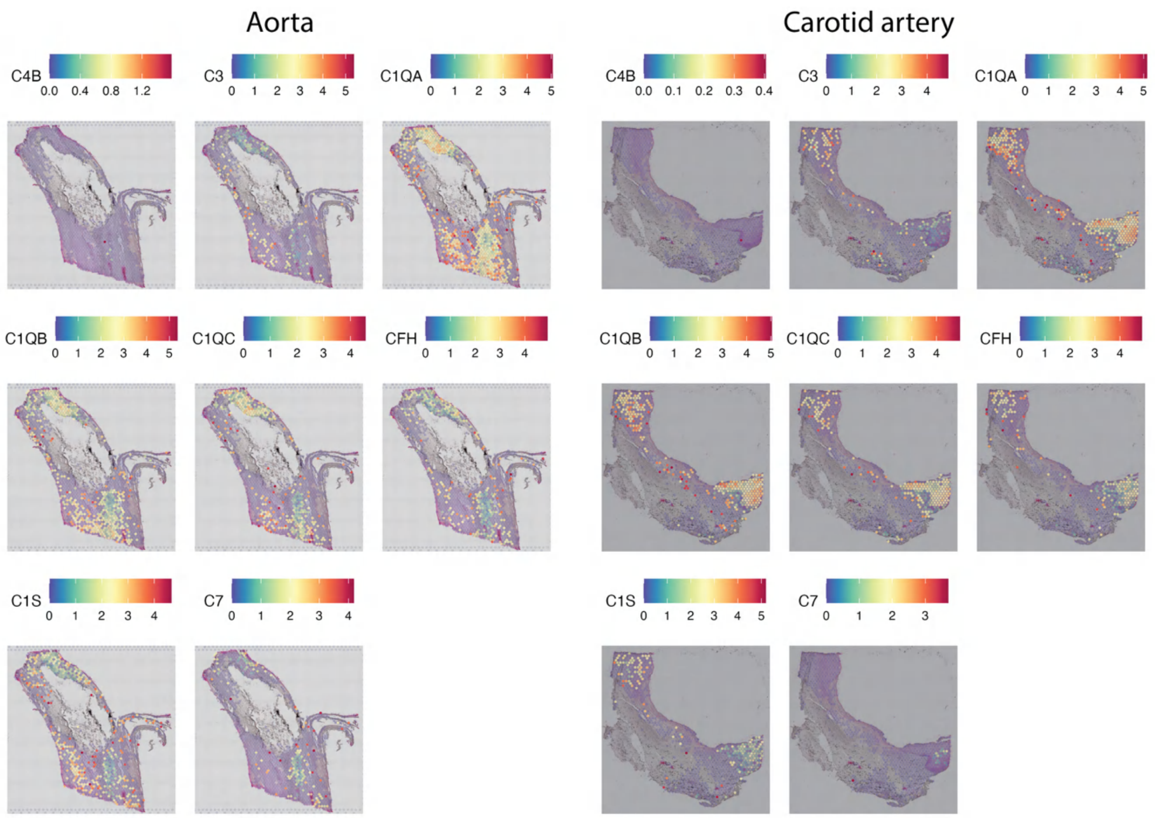


Supplementary Figure 11. Complement cascade component expression in aorta (three columns from the left) and carotid artery (three columns from the right). The colour bar shows normalized expression, where blue refers to no expression and red the maximum of the normalized expression, which varies between genes and samples. The active area on the Visium slides is 6.5x6.5 mm and each spot is 55 µm in diameter.

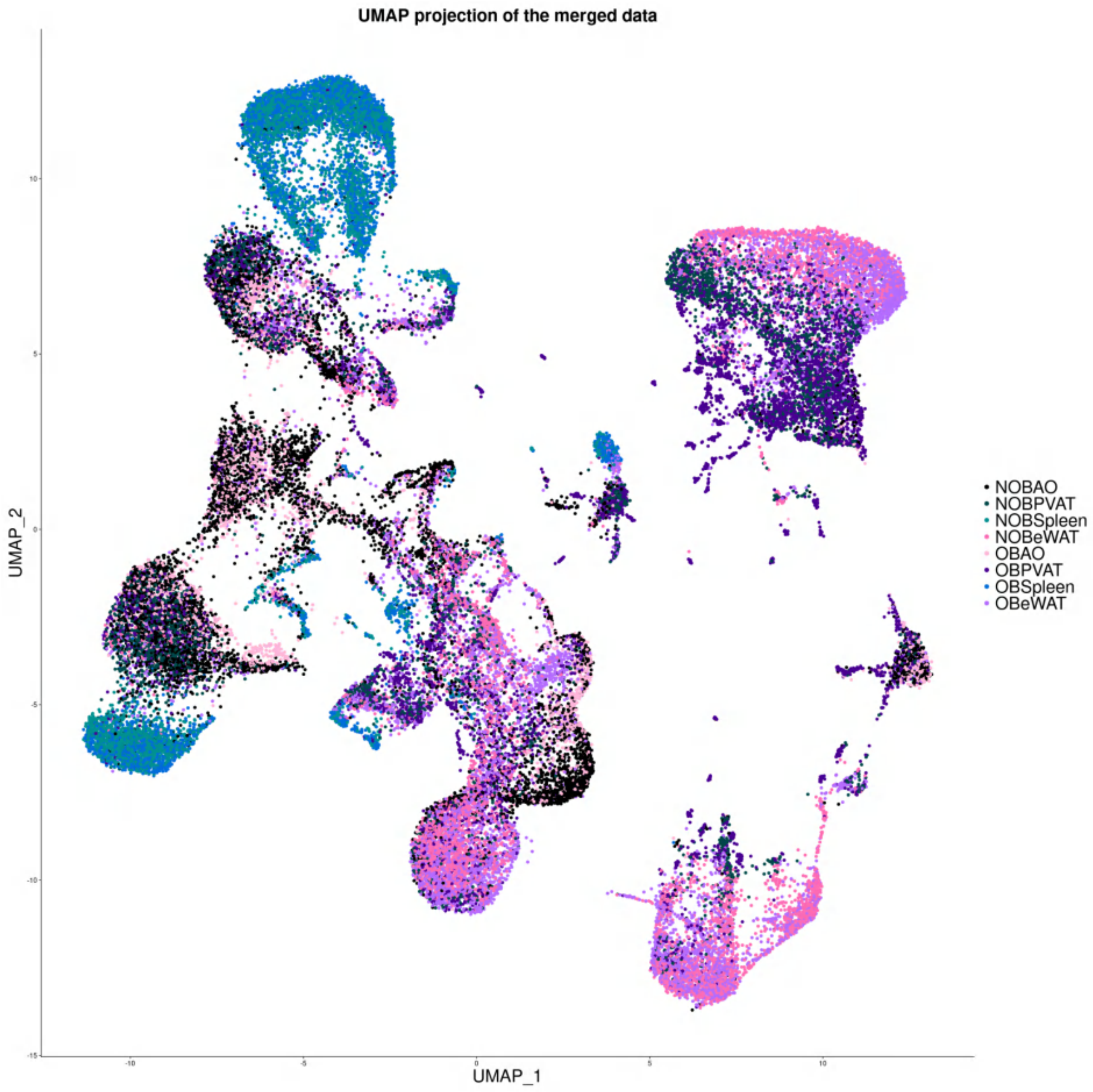


Supplementary Figure 12. UMAP projection of the merged data before integration with Harmony. Number of cells 47495.

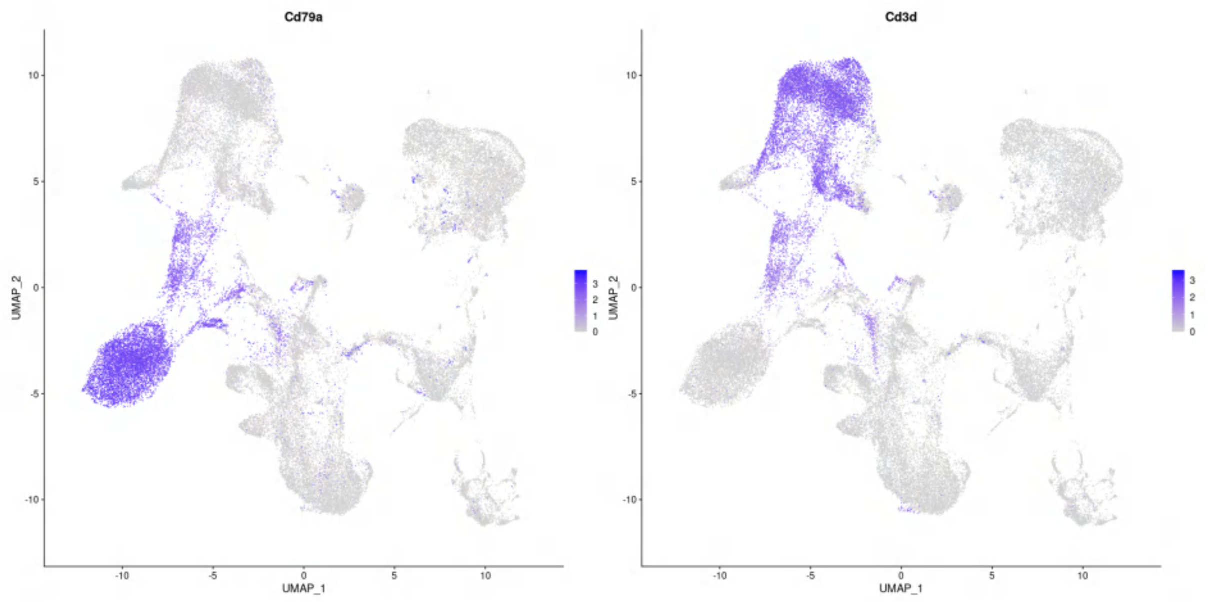


Supplementary Figure 13. *Cd79a* (left) and *Cd3d* (right) expression in feature plots showing a double positive cell population between the distinct *Cd79a+* (round population in the left lower corner) and *Cd3d+* (larger population in the right upper corner). These cells were removed from downstream analysis. Number of cells is 47495 in both pictures.

**Supplementary tables are provided separately as excel files.**

Supplementary Table 1. Details of antibodies for IHC and mIF validations in mouse PVAT-aorta tissues.

Supplementary Table 2. Differentially expressed genes between obese and non-obese states in aorta *Cd8+* T memory and dendritic cells.

Supplementary Table 3. Differentially expressed genes between obese and non-obese states in PVAT *Mgp+* and *Pi16+* fibroblasts, endothelial cells and intermediate monocytes.

Supplementary Table 4. Differentially expressed genes between obese and non-obese states in eWAT *Mgp+* fibroblasts, *Trem2+* macrophages and VSMCs.

Supplementary Table 5. Gene set enrichment analysis results of the up- and downregulated DE genes in PVAT-derived *Pi16+* and *Mgp+* fibroblasts.

Supplementary Table 6. Gene set enrichment analysis results of the up- and downregulated DE genes in eWAT-derived *Pi16+* and *Mgp+* fibroblasts.

### Methods

*Sample collection and processing for single-cell transcriptomics*

A total of ten male mice lacking Ldlr and solely expressing ApoB100 (Ldlr$^{-/-}$ / ApoB$^{100/100}$, C57BL/6J background; strain 003000, Jackson Laboratory) were bred and housed at the University of Turku Central Animal Laboratory (Turku, Finland). It is well established that Ldlr$^{-/-}$ / ApoB$^{100/100}$animals develop extensive atherosclerosis due to the markedly high abundance of small apoB100-containing lipoprotein particles[1]. This model accurately reflects human lipoprotein metabolism, which physiologically resembles LDLR deficiency more closely than ApoE deficiency.

Mice fed with high fat diet (TD.88137, Envigo, US) for 3.5 months represented obese or atherosclerotic model (n=5) and mice fed with regular chow diet represented non-obese state or controls (n=5). The mice were age matched at the time of the sacrifice (age ranging between 27 and 34 weeks). The mice were first anesthetized by isoflurane (induction 4-5 %, maintenance 2-2.5 %), followed by drawing of the blood with cardiac puncture, after which the animals were euthanized with cervical dislocation. Each mouse was perfused with ice-cold 10 ml DPBS (L0615-500, Biowest, Nuaillé, France) including 10 U/ml heparin (C29859, LEO Pharma, Ballerup, Denmark) before further dissection.

Four different tissues were collected: aorta (incl. the aortic tissues from ascending aorta until the renal arteries branch from the abdominal aorta), perivascular adipose tissue surrounding the aorta, epididymal white adipose tissue, and spleen. The tissues were kept in DPBS on ice, they were individually weighed, and PVAT was separated from aortas on a cold block in a drop of DPBS. Single-cell solutions were prepared from the four tissues by following the protocol specified in Figshare (10.6084/m9.figshare.29312777), including red blood cell lysis with ice-cold lysis buffer (eBioscience™ 10X RBC Lysis Buffer, 00-4300-54, Thermo Fisher Scientific, US). The same tissue types from all mice in the same group were pooled together from the beginning, except the aortas. The input quantities were equalized (for mass) to ensure similar proportions of each individual sample in the pooled sample. The aorta samples were pooled after dissociation. The cell concentration and viability of the resulting cell suspensions were measured with a LUNA FX7 instrument (Logos Biosystems, South Korea) using acridine orange and propidium iodide stains. The viability varied between ~ 87-95 %, therefore no dead cell removal was needed for any of the samples.

A positive enrichment of CD45+ cells was carried out using a Miltenyi CD45 MicroBeads kit (mouse, 130-052-301) in the pooled aorta sample. The enrichment was done according to the Miltenyi protocol (140-003-515.01). Before enrichment 30 µl of the original sample was taken aside so it could be mixed with the enriched sample. This was done because the enriched sample is expected to be highly pure (> 90 % CD45+ cells) and we did not want to exclude all the CD45- cell types. A 20-25%:75-80% ratio was used for mixing the original and enriched sample, respectively. Next, the cells were stained with an experimental BioLegend Totalseq-C panel, containing 138 antibodies (10.6084/m9.figshare.29312825, file feature_bc_totseqC.csv), following the manufacturer's protocol (https://www.biolegend.com/en-us/protocols/totalseq-b-or-c-with-10x-feature-barcoding-technology) and continued to the single-cell immune profiling protocol.

For validation experiments we got ten more mice from the Lab Animal Centre at the University of Eastern Finland. The mice were transferred to the mouse facilities of the University of Turku Central Animal Laboratory, and five of them were put on a high fat diet (TD.88137, Ssniff Spezialdiäten GmbH, Germany) for 3,5 months, while the other five were sacrificed at 6-month-old after being solely fed with chow (Teklad 2018c, Envigo, US). The same tissues were collected from the mice as in the original experiment. The samples were snap frozen in ethanol dry ice slurry and stored in -150℃ for later use. Also, part of the samples were collected so that the PVAT was kept around the descending aorta. These aorta-PVAT samples (approximately 10 mm in length) were collected rostrally just above the renal arteries, and were individually fixed with 4 % paraformaldehyde, dehydrated in 70 % EtOH, and paraffin embedded for immunohistochemistry (IHC) and multiplexed immunofluorescence (mIF) experiments.

*TotalSeq antibody staining of cell surface proteins*

From each tissue type, approximately 2.5x10^5 cells were used for surface staining, to be used as input material for the subsequent single-cell analysis. The cells were first incubated with anti-mouse CD16/CD32 (Clone 2.4G2) (InVivoMAb, PN-BE0307) 1:100 in 1% BSA-PBS for 10 minutes on ice to block Fc receptors. The lyophilized TotalSeq-C mouse custom cocktail panel (BioLegend, PN-900002539) was reconstituted following manufacturer's instructions. After blocking of the Fc receptors, the cell suspension was incubated with 12.5 µl of the TotalSeq-C panel and 75 µl of 1% BSA-PBS on ice for 30 minutes. After the antibody incubation, the cells were washed first once with 0.04% BSA-PBS and then twice with 1% BSA-PBS, pelleting the cells by centrifugation (400 x *g*, 5 min, +4℃) and removing supernatant between the washes. The cells were finally resuspended in 1% BSA-PBS in concentrations of 300-1000 cells per µl.

*Single-cell sequencing and preprocessing of the sequencing data*

A total of eight samples were sequenced; tissue-specific pooled samples from obese and non-obese atherosclerosis states, stained with the TotalSeq-C panel. The sequencing libraries for gene expression and cell surface proteins were prepared used the 10X Genomics Chromium Next GEM Single Cell 5' Reagent Kits v2 Dual Index protocol, following manufacturer's instructions (CG000330 Rev A; kits PN-1000263, PN-1000190, PN-1000256, PN-1000215). In brief, approximately 17,000 cells from each sample were loaded onto Chromium Single Cell K chips, aiming to capture 10,000 single cells per sample. Single cells were partitioned into gel bead emulsions using a Chromium Controller (10x Genomics). After reverse transcription and cDNA barcoding, the emulsions were reversed and the subsequent steps were performed in bulk. Full-length cDNA was amplified with PCR (14 cycles) using a Veriti cycler (Applied Biosystems / Thermo Fisher).  The libraries for the cell surface protein antibodies were prepared from unfragmented molecules. The libraries were finally indexed and amplified by PCR (14 cycles for gene expression libraries and 8 cycles for the cell surface protein libraries). The quality controls throughout the process were done with a Bioanalyzer 2100 (Agilent Technologies, US). The sequencing was done at Novogene using NovaSeq6000 with S4 flow cell. The reads were paired-end 150 bp. The data were preprocessed using 10X Genomics Cell Ranger V6 multi in a CSC - IT Center for Science (Finland) high performance computing (HPC) system with the configuration files in Figshare 10.6084/m9.figshare.29312825.

*Quality control and sample integration*

The single-cell analyses were run in R[2]. Scripts used for all the following analysis workflows are available from https://github.com/BioTechnoAlchemist/TERVA. The initial quality controls and normalization of individual sample data were run in R (v4.1.2) on a local computer (MacOS Catalina 10.15.5), but the subsequent integrated analyses were run on a CSC HPC interactive environment with R (v4.2.1), under the Red Hat Enterprise Linux 8. Our analysis workflow started with quality control processes: first ambient RNA removal with SoupX v1.5.2[3], followed by crude pruning of the data in Seurat v4.1.1[4], and ultimately doublet removal with DoubletFinder v2.0.3[5].

Cell-free RNA or “ambient RNA” is a common source of noise in scRNAseq data and therefore a step to remove this noise was incorporated into the quality control workflow. We used either an automated estimation of ambient RNA or if that seemed too lenient (ambient RNA estimated to be less than 2 %), a reference list of genes (*Igkc*, *Ighg1*, *Ighm*, *Iglc2*, *Iglc3*, *Iglc1*, *Ighe*, *Ighg3*, *Ighd*) was used to estimate the global contamination fraction.

Droplet-based snRNAseq data may contain empty droplets and/or multiplets, so to remove clear outliers a crude pruning of the data was done by setting thresholds where the number of features for a cell must be between 200 and 5000. In addition, it was determined that the mitochondrial content of a cell must be less than 5-10 % to pass the quality control (depending on the sample, which was evaluated visually from violin plot before and after applying different thresholds). After the pruning, a basic Seurat data pipeline was run with default parameters to normalize and scale the data, after which DoubletFinder was used to find and remove possible remaining multiplets based on the expected multiplet rate dependent of the cells loaded reported by 10X Genomics in the CG000330 Rev A manual.

After the quality control steps were done on each sample, the data was transferred to the CSC Puhti server, where it was merged. At this point genes *Gm42418* and *AY036118* were removed from the data as they are commonly causing technical noise[6], after which the data was renormalized and - scaled with the default parameters, followed by finding variable features using the “vst” method and number of the top variable features set to 3000, and principal component analysis (PCA) with the default parameters. Twenty first PC’s were chosen based on an elbow plot of the proportion of variance explained by the PC’s. Then uniform manifold approximation and projection (UMAP) dimensional reduction was run based on the PCA with 1:20 dimensions, followed by finding k-nearest neighbors (also using 1:20 dimensions), and finding clusters with resolution 0.6. The RunUMAP and RunPCA functions have a seed set to 42 by default and the same seed was used for the FindClusters function. The UMAP visualization of the merged data was inspected before data integration (Supplementary Fig 12). At this point one antibody was removed from the CITE-seq data because BioLegend reported a quality issue and mistaken identity in one antibody (Cd11a; correctly Cd28, clone 37.51), after which the CITE-seq data was also normalized using the centered log-ratio method. Moreover, because CITE-seq data may suffer from major background noise, to address this issue, we leveraged “dsb” (denoised and scaled by background) R package to normalize and denoise the protein expression data[7]. We defined the cell protein matrix and empty droplet matrix using the raw data produced by the CellRanger. To normalize the protein expression data, “DSBNormalizeProtein” function was used with “denoise.counts” and “use.isotype.control” parameters set to “TRUE”. "IgG1_kappa_isotypeCtrl", "IgG2a_kappa_isotypeCtrl", and "IgG2b_unknown_isotypeCtrl" were used as isotype controls. Each sample was processed separately and then integrated.

Next, the data was integrated with Harmony [8] to correct for any batch effects due to sample processing. We used “sample” as the variable to integrate out. Then the UMAP dimensional reduction technique was rerun with the harmony embeddings using the first twenty embeddings. Finding nearest neighbors and UMAP were also rerun with the same first twenty harmony embeddings. Then finding clusters was rerun with resolutions from 0 to 1.5 with 0.1 increments and Clustree-package (v0.5.0,[9]) was utilized to visually inspect an appropriate clustering resolution, which was evaluated to be 0.7. While visually inspecting some major immune cell markers with the FeaturePlot function, we found a cell population that was double positive for *Cd79a* and *Cd3d* genes (Supplementary Fig 13). Closer inspection of RNA expression in these cells showed that 1292/1555 (83.09 %) of the cells were concurrently positive for multiple B and T cell specific markers *Cd79a*, *Cd19*, *Cd3d*, *Cd3e* and *Cd3g*, and and 1380/1555 (88.75 %) were double-positive for the Cd19 and Cd3 proteins in the CITE-seq data. The majority of these multipositive cells were from the aorta samples (1374/1555 cells; 977 cells from the non-obese and 397 cells from the obese-derived samples) and because we could not exclude the possibility of technical causes for the multipositivity from the sample preparation and processing, this cluster was removed from the subsequent analyses, reducing the total cell count from 47495 to 45940.

*Cell annotation and differential gene expression analysis*

Cell types were defined by combining automated (SingleR v1.6.1 and celldex v1.6.0 packages [10]) and manual cell annotation aided by the CITE-seq data. We used MouseRNAseqData from celldex to create the reference for the automated annotation[11]. The subsequent differential gene expression analyses (between clusters for helping annotation and between atherosclerosis states in different cell types within tissues) were run with Wilcoxon rank sum test. Threshold for significance was adjusted P-value < 0.05.

*Gene set enrichment analysis*

Gene set enrichment analyses (GSEA) using GO, KEGG and Reactome databases were carried out with g:Profiler [12] (version *e110_eg57_p18_4e6dcbc3*) using their web interface at https://biit.cs.ut.ee/gprofiler/gost. The visualization was carried out on the g:profiler generated output files using the R-packages multienrichjam (0.0.82.900), enrichplot (v1.23.1.992) in a local environment (macOS Monterey 12.6.7, R-version 4.3.3). We used our own R-script to mine immune-related terms and pathways from the g:profiler output csv-files. The words used in this analysis were ("immune", "immunity", "inflammation", "inflammatory", "mhc", "complement", "c1", "c2", "c3", "c4", "hypersensitivity", "autoimmune", "leukocyte", "defense", "cytokine", "interferon", "interleukin", "antigen", "neutrophil", "granulocyte", "platelet", "macrophage", "th1", "th2", "th17", "dendritic cell", "t cell", "b cell", "rheuma", "arthritis", "atherosclerosis", "lupus", "diabetes", "viral", "bacterial", "graft").

*Trajectory analysis*

The trajectory analyses were run with the R-package Totem (v0.99.2)[13], which uses SingleCellExperiment-objects, and utilizes Slingshot[14]. First, the conversion from seurat-object to a SingleCellExperiment-object was done with the SingleCellExperiment R-package (v1.18.0) and the required dimensional reduction was carried out with the dyndimred R-package (v.1.0.4).

*Single-cell regulatory network inference and clustering (SCENIC) analysis*

We employed the pySCENIC (v0.12.1) Python package to infer transcription factors (TFs) within various fibroblast subpopulations[15]. Initially, we utilized GRNBoost2 to analyze Gene Regulatory Networks (GRNs) based on the co-expression patterns of TFs and their target genes. Next, we applied the cisTarget function to identify putative regulatory regions of the target genes. This function enabled the identification of enriched TFs within each module, where a module is defined as a TF and its potential target genes. The relative biological activity of the identified modules in individual cells was quantified using the AUCell function. Regulon Specificity Scores (RSS) were calculated using the Jensen–Shannon Divergence[16] to evaluate the cell-type specificity of the modules.

*Cell interaction analysis*

Network analyses were executed with the NicheNet package v1.1.1[17] in the CSC Puhti HPC environment. The required network and ligand-target matrix input files for the basic seurat-based NicheNet-pipeline were downloaded from https://zenodo.org/record/3260758.

The sender cell types were: B cells, *Mfap4+* fibroblasts, *Ccl11+* fibroblasts, *Mgp+* activated fibroblasts, *Lef1+ Tcf7+ Cd4+* T cells, *Gpihbp1+ Fabp4+* endothelial cells, classical and non-classical monocytes, conventional dendritic cells, *Folr2+ Lyve1+* M2 macrophages, *Ccl4+ Cxcl2+ Ccl3+* inflammatory macrophages, *Pf4+ Retnla+* macrophages, mesothelial cells, *Fscn1+ Apol7c+* dendritic cells, intermediate monocytes, NK cells, plasma cells, *s100a9+/a8+* granulocytes, and VSMCs.

*Sample collection of human carotid arteries and aorta for spatial transcriptomics*

A sample of atherosclerotic aorta was obtained from a 67-year-old male undergoing aorto-femoral bypass surgery due to chronic, symptomatic peripheral artery disease. Carotid artery plaque samples were obtained from a 61-year-old female and a 60-year-old male undergoing surgical endarterectomy due to critical carotid artery stenosis, with symptom onset less than two weeks. The study conforms to the Declaration of Helsinki, the institutional review boards of the Hospital District of Southwest Finland and Turku University Hospital approved the study (permit number T03/024/19, ethical statement number T75_2011), and patients gave written informed consent.

*Histological and immunofluorescence staining of human aorta and carotid artery sections*

The tissues were embedded in optimal cutting temperature compound, frozen at −70°C, and cut into 6 μm cryosections and stained with hematoxylin-eosin (H&E). The stained slides were scanned with a digital slide scanner (Pannoramic 1000 Flash; 3DHISTECH, Ltd., Budapest, Hungary). To examine the expression of CD68, Mannose receptor and Folate receptor-beta (FR-β; alias CD206) positive macrophages on the lesion region of the tissues. Subsequent adjacent sections were formalin fixed for 10 minutes then treated with epitope retrieval solution pH 6, Leica RE7113, followed by blocking with Normal antibody diluent (BD09-125, WellMed BV, Duiven, The Netherlands). These sections were double stained with mouse anti-human CD68 (1: 1000; catalog number: M0876, Dako Agilent, Glostrup, Denmark) and rabbit polyclonal anti-Mannose receptor antibody (1: 1000; catalog number: ab64693, Abcam, Cambridge, United Kingdom) followed by goat anti-Mouse Alexa Fluor 488 conjugated secondary antibody (1:1000; catalog number: A-11017, Invitrogen, ThermoFisher Scientific, Waltham, United States) and donkey anti-rabbit Alexa 549 (1:1000; catalog number: A21207, Invitrogen, ThermoFisher Scientific, Waltham, United States) or anti-human CD68 and anti-human FR-β allophycocyanin (APC)-conjugated antibody (1:100, mouse IgG2a; BioLegend, San Diego, CA, United States) followed by goat anti-Mouse Alexa Fluor 488 conjugated secondary antibody (1:1000; catalog number: A-11017, Invitrogen, ThermoFisher Scientific). Then the sections were mounted with ProLong Gold antifade with DAPI (catalog number: P36931, Invitrogen, ThermoFisher Scientific, Waltham, United States). The stained slides were scanned with a digital slide scanner (Pannoramic Midi fluorescence slide scanner; 3DHISTECH, Ltd., Budapest, Hungary), then images were made using Caseviewer software (3DHISTECH, Ltd., Budapest, Hungary). The H&E and IF staining were used to select the regions of interest for the spatial transcriptomics experiment.

*Spatial transcriptomics sample processing and sequencing*

A spatial transcriptomics experiment was carried out on human aorta and carotid artery sections using the 10X Genomics Visium (V1) gene expression platform. Of each sample, two technical replicates were generated from adjacent sections. Therefore, we carried out the spatial transcriptomics on a total of eight sections. The imaging required for the protocol was carried out by the researchers at the Cell Imaging and Cytometry core facility using a Nikon

Eclipse Ti2-E microscope equipped with a Nikon DS-Fi3 CMOS camera with the specifications explained in the CG000241 Visium Imaging Guidelines Rev B. RNA integrity number (RIN) was determined for the samples with the Bioanalyzer 2100 and Eukaryote Total RNA 6000 Pico kit (5067-1513, Agilent). The carotid arteries had RIN values of 6.10 and 7.70, and the aorta sample had a RIN value of 6.90. Optimal permeabilization time was determined from carotid samples and due to similar tissue structure, the same permeabilization time (18 minutes) was used for the aorta and carotid arteries. The protocols used were: Demonstrated Protocol Methanol Fixation and HE Staining CG000160 Rev C, Visium Spatial Tissue Optimization User Guide CG000238 Rev D, and Demonstrated Protocol Visium Spatial Protocols Tissue Preparation Guide CG000240 Rev D.

The sequencing libraries were constructed following the Visium Spatial Gene Expression User Guide CG000239 Rev F and cDNA and subsequent library quality was checked with the Bioanalyzer 2100 using the High Sensitivity DNA Assay (5067-4626, Agilent). The sequencing was carried out with Illumina NovaSeq using an SP flow cell with the recommended 28+10+10+90 bp read lengths, by the Finnish Functional Genomics Centre, and they used Space Ranger (v1.3.1) to generate the count matrices for downstream analyses. Space Ranger raised a low fraction reads in spots warning for the sections of the first region of the aorta samples, which may be a result of nonoptimal permeabilization. The other two aorta (second region) and four carotid artery sections had no flags. Subsequent QC was carried out separately for each section in Seurat (v5.3.0) in R (v4.4.1) in a local environment using an MacOS Sequoia v15.4.1 system. Low quality spots were filtered with variable threshold for number of genes per spot per section (40-90 for aorta and 60-100 for carotid artery samples). The thresholds for the mitochondrial and ribosomal gene content were 15 and 20 percent for each section, respectively. Before gene mapping, data log-normalization, scaling, and finding of variable features were carried out similar to the single-cell workflow.

*Immunohistochemistry and multiplexed immunofluorescence staining of mouse tissues*

IHC and multiplex-IF experiments were carried out on horizontal sections of the ten individual aorta-PVAT samples. The paraffin embedding and subsequent IHC/mIF stainings of the sections were carried out at the Histology core facility of the Institute of Biomedicine, University of Turku, Finland. For routine IHC, each experiment slide had a total of five stained sections so that each slide contained either two early disease and three late disease samples or vice versa.

For routine IHC, antigen retrieval for PI16 and CD74 was performed with a pressure cooker (Decloaking chamber NxGen, Biocare Medical) for 20 minutes using citrate buffer with pH 6.0 (BioSite, BSC-OKHURL) and for CD19 with a microwave (7 minutes 600 W + 7 min 450 W + 20 minutes of cooling down) with Tris-EDTA buffer with pH 9.0 (BioSite, BSC-2OIB1M). Staining was done with a semiautomated Labvision autostainer (Thermo Fisher Scientific). Washing buffer was 0.05 M Tris-HCL with pH 7.6 (Reagena, 112270) and 0.05 % Tween 20 as a detergent. Blocking was carried out with hydrogen peroxide and Normal antibody diluent (BD09, Immunologic) or Dako REAL antibody diluent (S2022, Agilent) (see * in Supplementary

Table 1) in room temperature (RT). The primary and secondary antibodies are listed in Supplementary Table 1. PI16, CD74, and CD19 primary antibodies were incubated for 60 minutes in RT, followed by secondary antibody incubation for 30 minutes. BrightDAB substrate (WellMed, BS04-110) was used with 10-minute incubation, followed by counter staining with Mayer's hematoxylin (1 minute in RT).

Multiplex-IF was performed by using Leica BOND RX automated stainer (Leica Biosystems). After 30 minutes of baking at 60°C and paraffin removal (Leica BOND Dewax Solution, AR9222, Leica Biosystems), slides were heated for 20 minutes at 100°C in antigen retrieval solution (Leica BOND Epitope Retrieval Solution 2, AR940, Leica Biosystems) and blocked for endogenous peroxidases in 3% hydrogen peroxide for 10 minutes at RT. Total of four staining rounds were carried out as follows: Protein blocking by Normal antibody diluent (10 minutes at RT) or Dako REAL antibody diluent (for PI16), primary antibody incubation (20 minutes at RT, Supplementary Table 1), secondary antibody incubation (10 minutes at RT, Supplementary Table 1), tyramide signal amplification by incubating slides with corresponding fluorophore-conjugated tyramide (Supplementary Table 1) for 10 minutes in 2 nM final concentration in Tyramide Amplification Buffer Plus (15129, Biotium), supplemented with fresh 0.0015% hydrogen peroxide. Antibodies were removed by heating slides for 10 minutes at 98°C in antigen retrieval solution (Leica BOND Epitope Retrieval Solution 2) and the whole staining cycle was repeated for each marker in the following order: PI16, F4/80, CD4, CD3. Slides were washed with Bond Wash Solution (AR9590, Leica Biosystems) in between steps and mounted with ProLong Gold Antifade Mountant with DAPI (P36935, Thermo Fisher Scientific). As negative control for F4/80, CD3e and CD4, rabbit IgG isotype control was used instead of primary antibody (Normal antibody diluent for PI16 as negative control). Imaging of the mIF-stained tissues was done with Leica Thunder widefield fluorescence microscope at the Cell Imaging and Cytometry core facility, at Turku Bioscience Centre.

*Tissue imaging and image analyses (PI16+ cell counts and morphometric analysis)*

Imaging of all mouse tissues was carried out by the researchers at the Cell Imaging and Cytometry core facility using a Nikon Eclipse Ti2-E microscope equipped with a Nikon DS-Fi3 CMOS camera and/or Pannoramic P1000 slide scanner (3DHISTECH, Budapest, Hungary). Image annotations for the subsequent image analysis were done with QuPath[18].

The morphometric analysis was carried out at the Medisiina Imaging Centre, University of Turku. The quantitative analysis of the scanned histological sections was performed using Visiopharm software version 2022.07 (Horsholm, Denmark). First, adipose tissue areas were labelled by a deep learning classifier, and the predicted labels were corrected manually where necessary. Adipose cells were then labelled using a workflow adapted from Iron Hematoxylin, Adipose Tissue analysis protocol package (#10113, visiopharm.com/app-center). Parameters of the original analysis protocol package were adjusted to optimize the correct detection of the adipose cells in the current tissues and staining. Areas of labelled adipose cells were extracted as the output. The data acquired from the previous steps was then analysed locally using a MacOS Sequoia v15.4.1 with the R (v4.4.1) base package stats, dplyr (v1.1.4) and visualized with ggplot2 (v3.5.2). The normality of the data was assessed using Shapiro-Wilk test, and after determining that the data is drawn from a normal distribution, we decided to use a one-way (two-sample) Welch t-test.

For counting of the PI16+ fibroblasts, we used the nucleus detection method from[19] and fine-tuned it for IHC staining data by applying domain adaptations (DA) in two steps. For the first DA step, positive detections were used as pseudo labels (both detection thresholds set to 0.4) and 1000 detections were randomly chosen for re-training the network (5 epochs). Second DA step was then performed using detections by the network from previous training round (thresholds 0.6 and 0.2), this time 10000 randomly selected detections were used for training (50 epochs). The resulting network was then used for nucleus detection form regions of interest. Positively stained cells were defined by thresholding (0.2) the brown channel of detected nuclei locations.

The following statistical analysis to compare estimated mean cell counts of obese and non-obese groups was carried out locally (as described above) in R. First, to examine the group differences we fit a generalized linear model using beta distribution with glmmTMB- package (v1.1.11)[20], which is an appropriate strategy for analysing proportions that can only vary between 0-1. Then we evaluated the model fit using DHARMa (v0.4.7)[21]. To compare the estimates from the generalized linear model, a Type III Anova test was done with the car-package (v3.1-3)[22]. Finally, we used emmeans-package (v1.11.1)[23] to generate the estimated marginal means, which were used to inspect how the estimated group means diverged. The emmeans-package was also used to visualize the results.
